# Mammalian aging involves genome-wide splicing degeneration leading to functional decline

**DOI:** 10.64898/2026.06.26.734787

**Authors:** Sirui Zhang, Alexander Tyshkovskiy, Kejun Ying, Siqi Wang, Vadim N. Gladyshev

**Affiliations:** Division of Genetics, Department of Medicine, Brigham and Women’s Hospital, Harvard Medical School, Boston, MA 02115, USA; CAS Key Laboratory of Computational Biology, Shanghai Institute of Nutrition and Health, University of Chinese Academy of Sciences, Chinese Academy of Sciences, Shanghai 200031, China

## Abstract

Alternative splicing exhibits significant changes during development and aging, affecting the composition and variance in the transcriptome. However, it is unclear whether and how age-associated splicing dysregulation leads to functional consequences. Here, an integrative analysis of transcriptome data across mouse and human tissues revealed that aging is characterized by systematic deterioration of the fidelity of RNA splicing, here termed splicing degeneration, a measure of functional alteration of reading frame and domain configuration of protein products. Genes with higher aging-associated splicing degeneration were more conserved and enriched for processes such as RNA metabolism and antigen presentation. By assessing alternative splicing events associated with functional deterioration, we quantified the degree of splicing degeneration. Its level increased with age but was alleviated following calorie restriction or rapamycin treatment, indicating that it can serve as a new molecular hallmark of aging. Mechanistically, through a comprehensive meta-data analysis, we discovered that splicing degeneration is associated with age-associated changes in specific splicing factors, which in turn showed a strong association with age-related transcriptome changes. Overall, our study demonstrates the intricate relationship between aging and genome-wide splicing degeneration, revealing a promising target for aging interventions acting to reverse splicing degeneration.

## Introduction

Aging is a complex biological process characterized by the accumulation of molecular damage, gradual decline in the physiological function and increased susceptibility to disease ^1,2^. The aging process is also accompanied by various molecular hallmarks, such as telomere attrition, cell exhaustion, and mitochondrial dysfunction ^3^. At the cellular level, changes in many processes are thought to contribute to aging, e.g. gene expression, mutations, genome instability, and dysregulation of transcription and translation ^4–7^. In recent years, enormous effort has been invested to understand the underlying molecular mechanisms that cause aging and its associated physiological changes, with the goal of developing effective approaches to slow down the aging process. However, the intricate mechanisms of how the disruption in gene regulation leads to age-related functional decline remains unclear.

Alternative splicing (AS) of mRNA is a fundamental step in gene expression where the different exons of a single gene can be selectively combined, allowing a single gene to produce multiple protein isoforms ^8^.The majority (>90%) of all human genes undergo AS ^9^.The selective inclusion of specific exons through AS often produces protein isoforms that have distinct structures, functions, or subcellular localization. Therefore, dysregulation of AS was found to cause many human diseases ^10–12^. AS is regulated by many *trans*-acting splicing factors that specifically recognize *cis*-elements (splicing enhancers or silencers) in the pre-mRNAs ^13,14^. In addition, the AS regulation is affected by transcription initiation and elongation, as splicing of most exons occurs co-transcriptionally. As a result, the epigenetic regulation of gene expression, such as methylation of histones and DNA, also has a direct impact on AS ^15^.

Previous studies indicated that AS of specific genes plays important roles in development ^16,17^ and aging ^18,19^, with many AS events and splicing factors changing during the entire lifespan of an organism ^20,21^. For example, Kun et al. identified thousands of age-associated splicing events, revealing that the genome-wide splicing profile may serve as a predictor of biological age ^22^. Mariotti et. al also discovered the striking accumulation of intron retention during the aging process ^23^. In addition, genes encoding core spliceosomal components and regulatory splicing factors have been identified to have a close relationship with aging ^19,24^. For instance, regulation of SRSF1 is associated with cellular senescence, and changes in its expression affect cell lifespan ^25,26^. Furthermore, the expression of HNRNPA1 decreases during aging, which impacts the splicing of many genes and is associated with the progression of neurodegenerative diseases ^27^.

Since aging is associated with the systemic decline of gene functions, we were intrigued by the possibility of a systematic AS change during aging and the potential functional consequences of such change. While many age-related AS events have been identified, they may resemble adaptive responses to the accumulation of damage rather than detrimental effects. At the same time, the association between dysfunctional damaging splicing events and age has not been firmly established.

The availability of a large amount of RNA-seq data provides an opportunity to systematically examine aging-associated transcriptome changes at the level of distinct splicing isoforms. By analyzing thousands of transcriptome profiling data across human and mouse tissues, we found that aging is accompanied by deregulation of AS, which in turn is associated with increased noise in splicing regulation. This age-associated deregulation of AS, which we term splicing degeneration, can introduce premature stop codons (PTC) and alter the reading frame and/or domain configuration of protein products leading to functional deterioration. We discovered that aging-related splicing degeneration can be alleviated by several anti-aging interventions and may serve as a new molecular signature of aging. Mechanistically, we revealed a “self-regulation” mode of splicing, wherein the misregulation of splicing factors themselves can impact splicing degeneration. Finally, by combining RNA interference sequencing (RNAi-seq) and enhanced Cross-Linking and ImmunoPrecipitation sequencing (eCLIP-seq), we performed a comprehensive analysis of the regulatory role of splicing factors. We identified key SFs that regulate aging-related splicing degeneration and found that abnormalities in SFs strongly influence age-related transcriptome changes. Overall, our study reveals an intricate relationship between aging and genome-wide splicing changes, suggesting that degenerated splicing is an attractive target for anti-aging interventions.

## RESULTS

### Differential splicing increases with age in older tissues

To comprehensively map the landscape of alternative splicing changes during aging, we first focused on 16,627 RNA-seq samples derived from 56 tissues collected by the Genotype-Tissue Expression (GTEx) project. The AS levels of all genes in each sample were quantified using paean ^28^, from which a percent-spliced-in (PSI) value was calculated for each AS event (Figure 1A). Using the baseline of the splicing pattern observed in young samples (younger than 35 years old), we compared the PSI value for each AS event identified in each aged sample (older than 45 years) with the average PSI level of young samples. For each old sample, the AS events with |delta PSI value| > 0.1 and P < 0.05 were identified as differential AS events. We observed a progressive increase in the frequency of differential splicing events with advanced age (Figure 1B). This trend persisted in different types of AS events (Figure S1A) and across prominent aging-related genes and pathways such as mTOR, AMP-activated protein kinase (AMPK) signaling and sirtuins (SIRTs) (Figure S1B). To strengthen the validity of our results, we adjusted the cutoff criteria for defining the young and aged groups, and the trend remained consistent (Figure S2A). Additionally, we employed an alternative method to identify differential AS events, using |log₂FoldChange| > 1 and P < 0.05 (one-sample t-test) as thresholds for each aged sample and calculated the correlation of differential AS counts and age (Figure S2B). To further investigate, we also applied a differential expression approach to examine the differential AS events between each old group and young group using the criteria |log₂FoldChange| > 1 and FDR < 0.05 (Figure S2C). Collectively, these results consistently revealed an upsurge in differential splicing events throughout the aging process, highlighting the increased disruption of alternative splicing. Using PSI levels of AS events, we further constructed a novel splicing aging clock and found a strong correlation between the predicted age and the actual age in the validation set (the Pearson r is more than 0.6 in most tissues), further highlighting AS changes as a crucial feature of the aging process (Figure S3).

**Figure 1.**
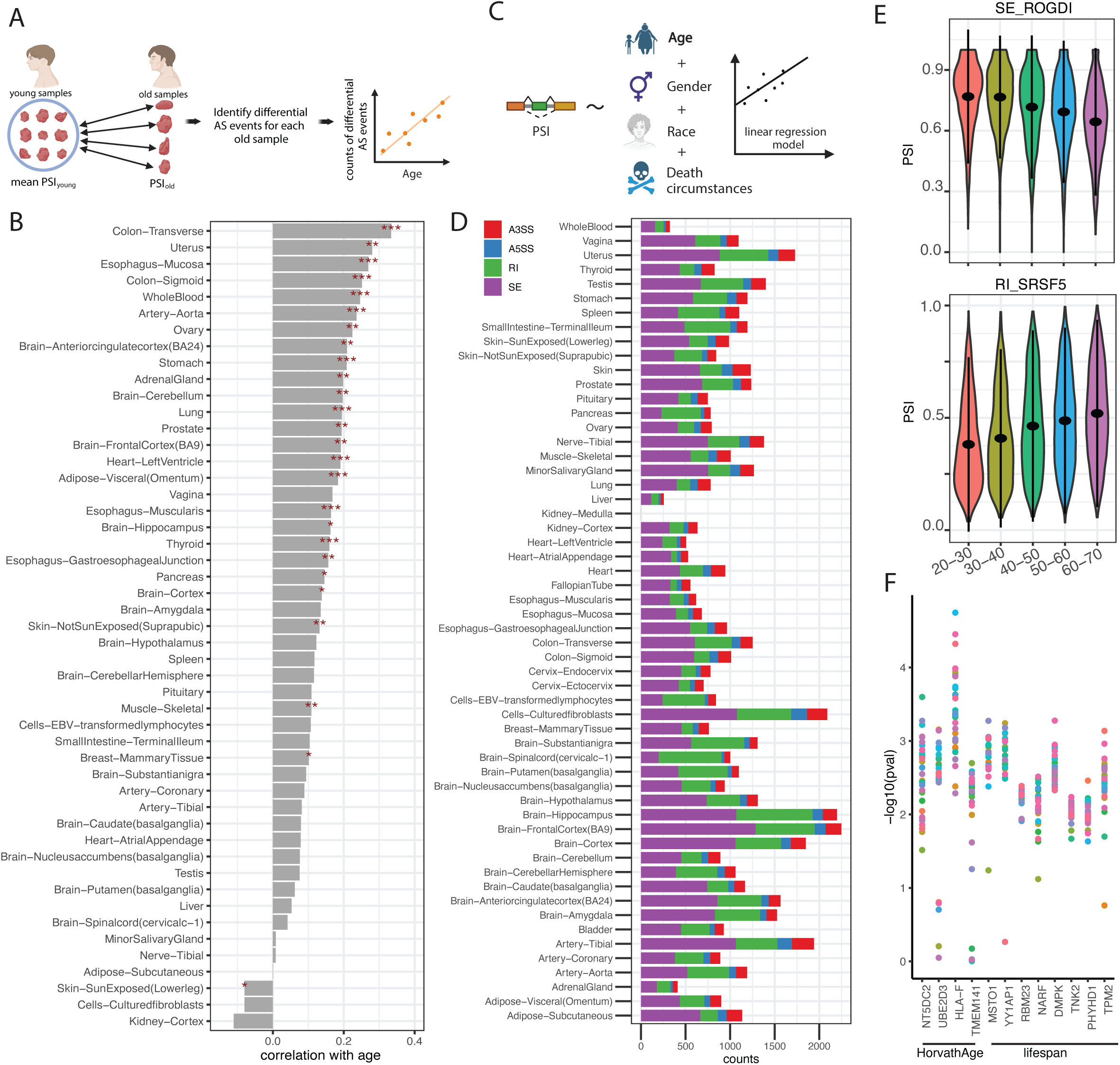
Differential AS events are increased with age. (A) The workflow to quantify differential AS in old tissues (older than 45 years) based on human data from GTEx. The PSI value for each AS event identified in each aged sample are compared with the average PSI level of young samples (younger than 35 years). For each old sample, the AS events with |delta PSI value| > 0.1 and P < 0.05 were identified as differential AS events. (B) Correlation of the number of differential AS events with age in indicated human tissues of GTEx. Asterisks indicate the significance of the Pearson correlation coefficient, *: P<0.05, **: P<0.01, ***: P<0.001. (C) Strategy for identifying aging-related AS events based on human tissue data from GTEx. (D) Counts of aging-related AS events in each tissue. Different colors indicate four types of AS. A3SS: alternative 3’ splice site, A5SS: alternative 5’ splice site, RI: retain intron, SE: skip exon. (E) Two examples of aging-related AS events identified from GTEx. The top is the PSI value distribution of the exon skipping of the fourth exon of *ROGDI* in different age groups, and the bottom example is the PSI value distribution of the intron retention event of the fifth intron of SRSF5 in different age groups. (***: P<0.001, ANOVA test). (F) The P values distribution of the causal effect (generated by GSMR) of aging-related AS on Horvath Age or lifespan in each tissue. The genes showing significant causal effects in at least 15 tissues are shown. The points in each column represent different tissues (also see Figure S6A).

### Aging-associated AS events may lead to functional consequences

To further investigate alterations in AS events associated with aging, we used a linear regression model to identify the AS events associated with age in each human tissue. With stringent criteria of (1) P value for β1 (coefficient of “AGE” in the linear regression model) < 0.05; (2) P value for Pearson correlation test between PSI value and age <0.05; and (3) β1 * Pearson correlation coefficient (between PSI values of specific AS events and age) >0 (Figure 1C, and Methods), hundreds of age-related AS events across diverse tissues were revealed (Figure 1D and S4). Notably, the aging process exhibited a significant increase in skipping the fourth exon of Rogdi Atypical Leucine Zipper (*ROGDI*), as well as in the retention of the fifth intron of Serine And Arginine Rich Splicing Factor 5 (*SRSF5*) (Figure 1E and S5). These alterations should have direct functional implications as exon skipping of *ROGDI* induces a frameshift, and intron retention of *SRSF5* generates to a premature termination codon (PTC).

We subsequently focused on the AS events correlated with aging across at least 10 tissues. In total, 1,509 such AS events were identified and termed aging-related AS events (Supplementary Table 1). To investigate causal relationships between AS and aging, we employed Generalized Summary-data-based Mendelian Randomisation (GSMR) using aging-associated AS events from GTEx, with the Horvath Age and lifespan as the outcomes ^29,30^. The GSMR assay measures how likely the change of certain genetic traits (splicing changes in our case) may cause the observed phenotype (the Horvath age or lifespan in our case). We found that multiple AS events exhibited significant causal effects. The genes with significant causal effects in at least 15 tissues were shown in Figure 1F as examples, and we also showed genes with significant causal effects in at least 5 tissues (Figure S6B-C). There are also several events show consistent results between the two outcomes (Figure S6D). All of these results suggest that many of the AS events identified may indeed play a causal role in aging-related phenotypical outcomes.

To further explore the relationship between age and AS, we investigated the common characteristics of aging-related AS events. First, we found that they exhibit a length bias, with the skipped exons being shorter and retained introns being longer (Figure 2A). Further analyses revealed a significant decrease in the 5’ splice site strength among aging-related AS events, while the 3’ splice site strength exhibited a lower, albeit non-significant trend compared to other AS events (Figure 2B). Notably, aging-related AS events were enriched in the coding sequence (CDS) and reduced in non-coding regions (Figure 2C), suggesting a potential significance of AS misregulation during aging. Additionally, these altered regions displayed higher conservation across 100 species compared to other AS events (Figure 2D), further supporting the notion that aging-related AS misregulation may impact the function of evolutionarily conserved genes.

**Figure 2.**
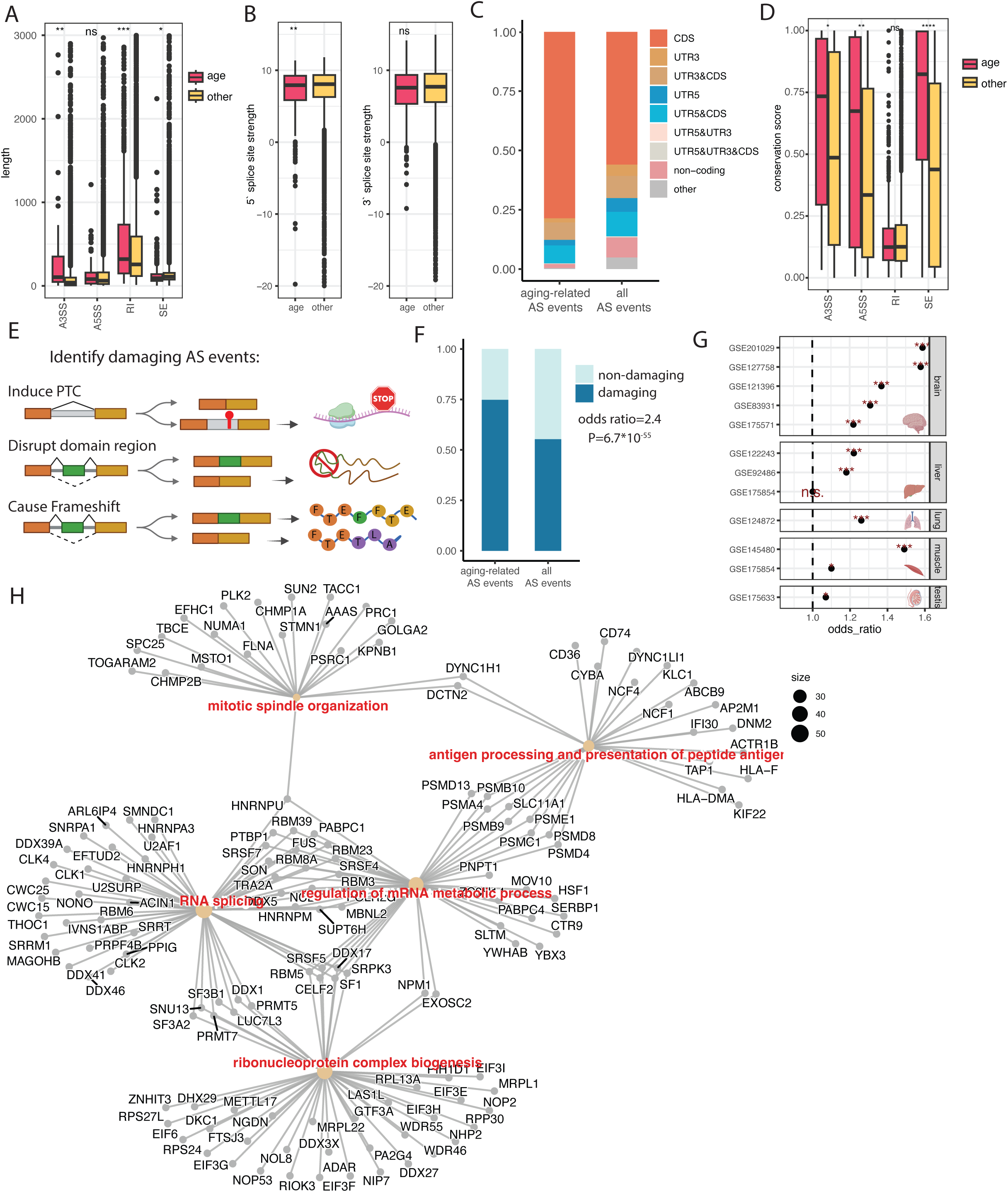
Features of aging-related AS events. (A) Length distribution of alternative forms (exon length in SE, intron length in IR, distance between differential splice sites in A3SS and A5SS) involving aging-related AS events (denoted as age) and all other AS events (other). (*: P<0.05, **:P<0.01, ***:P<0.001, Student’s t-test). A3SS: alternative 3’ splice site, A5SS: alternative 5’ splice site, RI: retain intron, SE: skip exon. (B) Distribution of splice site strength of aging-related AS events (age) and other AS (other) events. (**:P<0.01, Student’s t-test). (C) Proportions of aging-related AS events and all AS events in different gene and genome regions. (D) Distribution of the conservation score of aging-associated AS events and all AS events among 100 placental mammals. (*: P<0.05, **:P<0.01, ***:P<0.001, Student’s t-test). (E) The workflow to identify the AS events with damaging consequences. There are three types of such events. The upper type shows an intron retention event that induces a premature termination codon (PTC). The middle shows an exon skipping event with the peptide translated by this exon within a protein domain region. The bottom type shows the AS event in which the deletion of an exon (or part of an exon) causes a frameshift. Also see Methods for detail. (F) Proportions of aging-related AS events and all AS events with damaging consequences or non-damaging consequences. Odds ratio and P value were calculated by Fisher’s exact test. (G) Odds ratio of the enrichment of aging-related AS events with damaging consequences compared with all AS events in several mouse datasets (indicated by accession numbers in GEO). Odds ratios and P values were calculated by Fisher’s exact test. *: P<0.05, **: P<0.01, ***: P<0.001. (H) Gene ontology of genes harboring damaging aging-related AS events.

Given that certain AS events exert only a marginal impact, while others may significantly influence protein structure and function, we posit that the AS events associated with age are more likely to harbor functional consequences, based on the above results. To further study the functional consequences of AS changes during aging, we classified all AS events as “damaging” or “non-damaging”. AS events with damaging consequences were defined as the following three types: (1) inducing a premature termination codon (PTC); (2) causing frameshift, and (3) located in a protein domain region (Figure 2E, and Methods). It is worth mentioning that our classification is to distinguish whether one of the two isoforms produced by alternative splicing will affect gene function. We believe that damaging type of AS events could influence the protein function while nondamaging events exert marginal impacts. Intriguingly, we found that aging-related AS events were enriched in damaging AS events (Figure 2F), suggesting a substantial impact on protein function during aging. Consistent results were also observed by analyzing several individual datasets of mouse tissues (Figure 2G), underscoring the generalizability of these findings that are not specific to a specific species or specific dataset. Gene ontology analyses showed that the genes with functions in RNA metabolism, RNA splicing, and antigen processing were enriched in aging-related damaging AS events (Figure 2H and S7), emphasizing that the disorder of AS during aging has a broad impact on crucial biological processes, including splicing machinery itself.

### Aging is accompanied by degeneration of alternative splicing

Given the enrichment of AS events with damaging consequences during the aging process, we postulated that this type of damage accumulates in older samples. To validate this hypothesis, we first examined the overall trend of the PSI value changes during aging and revealed a decline in the mean PSI values for skipped exon (SE), alternative 3’ splice site (A3SS) and alternative 5’ splice site (A5SS), as well as an increase in the PSI values for intron retention (IR) (Figure S8A), which is similar to previous studies ^23^ and further suggest a functional decline of splicing during aging.

To further quantify splicing degeneration, we only considered the AS events with damaging consequences. Similar to the classification in Figure 2H, there are three types of damaging AS in total. First, for IR events with a possibility of inducing a PTC, introns that are retained during aging could affect protein expression and could be considered as damaging events. Second, for SE events with the exon located in a protein domain region, exon skipping during aging could affect protein function. For the SE, A3SS and A5SS events which cause a frameshift, in most (∼90%) genes, the long isoforms are the dominant isoforms at the protein level ^31^. Hence, it appears that shorter isoforms during aging are those that likely cause functional deterioration, i.e, those AS events that cause frameshift with low PSI values tend to be damaging. Based on this finding, we established a computational pipeline to quantify splicing degeneration levels by assessing the proportion of damaging isoforms (Figure 3A). In essence, we aimed to quantify the level of damaging SE, A3SS, and A5SS events with lower PSI values and damaging IR events with higher PSI values.

**Figure 3.**
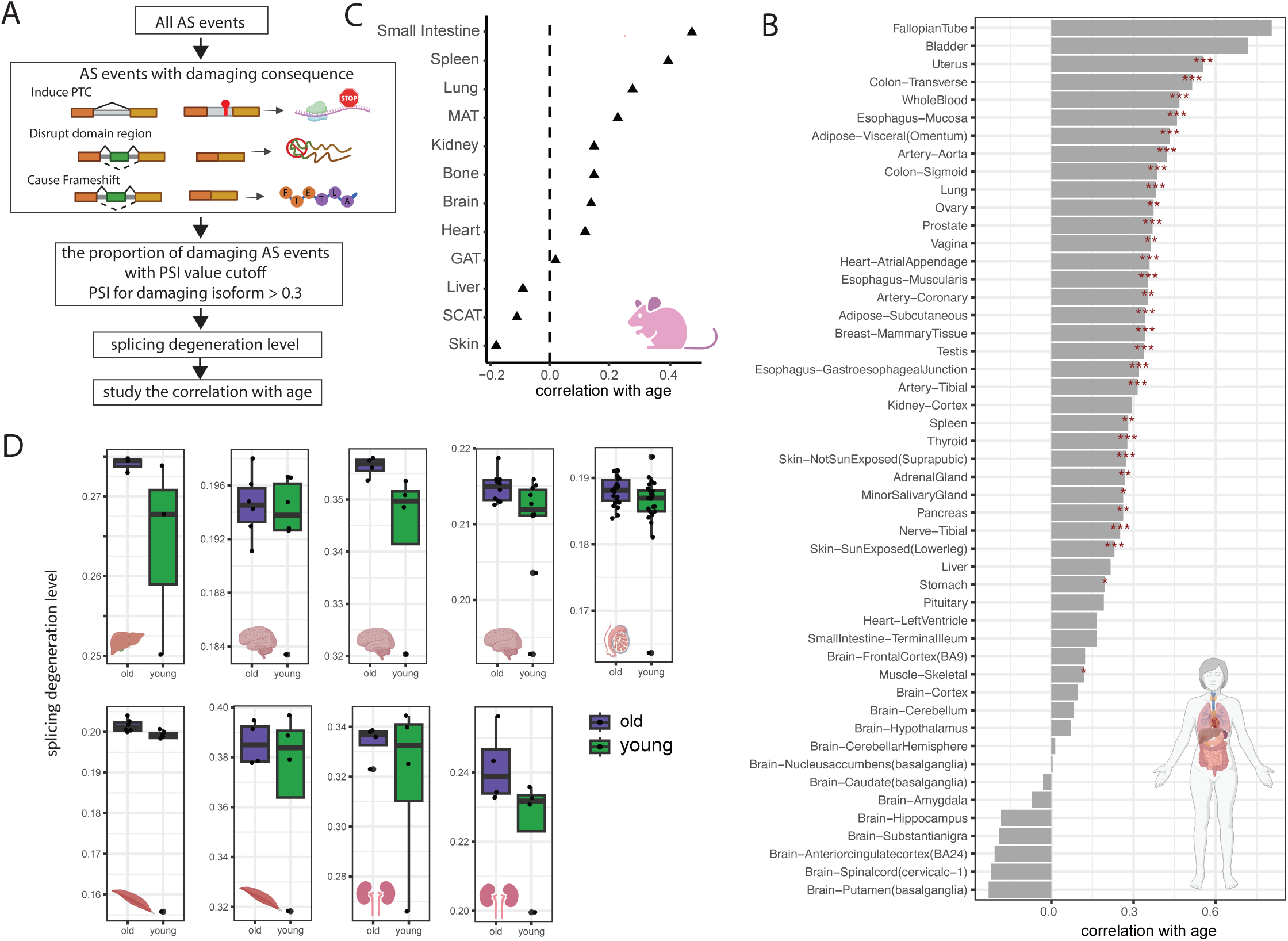
Aging is accompanied by degeneration of alternative splicing. (A) Workflow to quantify AS degeneration levels. Three categories of damaging AS events are counted. For SE, A3SS, and A5SS events, the proportion of PSI values less than 0.7 was counted, and for RI events, the proportion of events with PSI values greater than 0.3 was counted. Proportion of damaging AS events with the above PSI cutoff was defined as the splicing degeneration level. See Methods for detail. (B) Correlation between splicing degeneration level and age in each human tissue (data from GTEx). Asterisks indicate significance of the Pearson correlation, *: P<0.05, **: P<0.01, ***: P<0.001. (C) Correlation between splicing degeneration level and age in each mouse tissue (data from Tabula Muris Consortium). (D) Splicing degeneration level of young and old mouse samples in several datasets (data from GEO). Tissues analyzed were liver, brain, testis, muscle, and kidney as indicated. The P values are 0.29, 0.36, 0.10, 0.041, 0.066, 0.25, 0.21, 0.40, 0.08 (Student’s t-test).

Through testing different cutoffs (see Methods), we observed a positive correlation between splicing degeneration levels and age under most cutoffs (Figure S8B). Using the proportion of damaging SE, A3SS and A5SS events with the PSI values less than 0.7, along with the damaging IR events with PSI values above 0.3, we set up a score to quantify splicing degeneration levels (Figure 3A), which is significantly increased during aging in most tissues (Figure 3B). Since the type of PTC and frameshift are more deleterious, we also used only these two types to quantify the splicing degeneration level, and the results showed a consistent trend (Figure S8C). We also applied this pipeline to mouse datasets, including the data from Tabula Muris ^32^ and several other individual datasets from GEO, and the results showed that splicing degeneration levels are positively correlated with age in most datasets and most tissues (Figure 3C-D). In both human and mouse analyses, we found that certain tissues exhibited high correlation consistency. For example, in the lung, splicing degeneration was strongly correlated with age in both species and ranked among the highest across multiple tissues. Among the common tissues of both species, spleen and small intestine also showed relatively consistent correlations. Collectively, all of the above evidence suggests that aging is intricately linked to the degeneration of alternative splicing, marked by the reduction in functional isoforms. Intriguingly, we also found that the splicing degeneration level exhibits an elevation in tumor samples compared to adjacent normal tissues, hinting at potential shared disorder patterns in splicing between aging and cancer (Figure S8D, data from TCGA).

### Splicing degeneration can be reversed by longevity interventions

In order to investigate whether splicing degeneration is functionally linked to aging or merely a consequence of the passage of time, we explored the impact of longevity interventions on AS. We first focused on rapamycin, known for its anti-aging effects ^33,34^, and collected multiple datasets of human cells treated with rapamycin, analyzing the AS changes before and after treatment. The results showed that among eight datasets representing human cells subjected to rapamycin, seven exhibited a reduction in overall splicing degeneration levels following rapamycin treatment, and only one dataset showed no differences (Figure 4A). Consistently, everolimus, a rapamycin analog with documented immune-boosting effects in older individuals and potential anti-aging properties ^35,36^, also demonstrated a decrease in splicing degeneration levels in three out of four datasets in the treatment group (Figure 4B). To comprehensively validate the impact of rapamycin *in vivo*, we extended our analysis to include aged mice treated with rapamycin, revealing similar results wherein the splicing degeneration level is decreased in rapamycin-treated mice (Figure 4C), suggesting that splicing degeneration is associated with aging rather than with the passage of time. Given rapamycin’s inhibition of mTOR activity and the pivotal role of the mTOR signaling pathway in controlling U2AF protein ^37^ – the major splicing machinery, we speculate that restoration of alternative splicing is one of the mechanisms by which rapamycin delays aging. To delve deeper into the relationship between rapamycin treatment and AS, we identified a set of 599 AS events that changed in at least three datasets in response to rapamycin treatment. We found that the genes harboring rapamycin-associated AS events significantly overlapped with the genes harboring aging-related AS events (Figure S9A), and functionally enriched in cell cycle and histone modification-related pathways (Figure S9B). In most rapamycin datasets, there were fewer retained introns and skipped exons after treatment compared to control groups (Figure S9C), suggesting a potential mechanistic link between rapamycin and splicing degeneration regulation.

**Figure 4.**
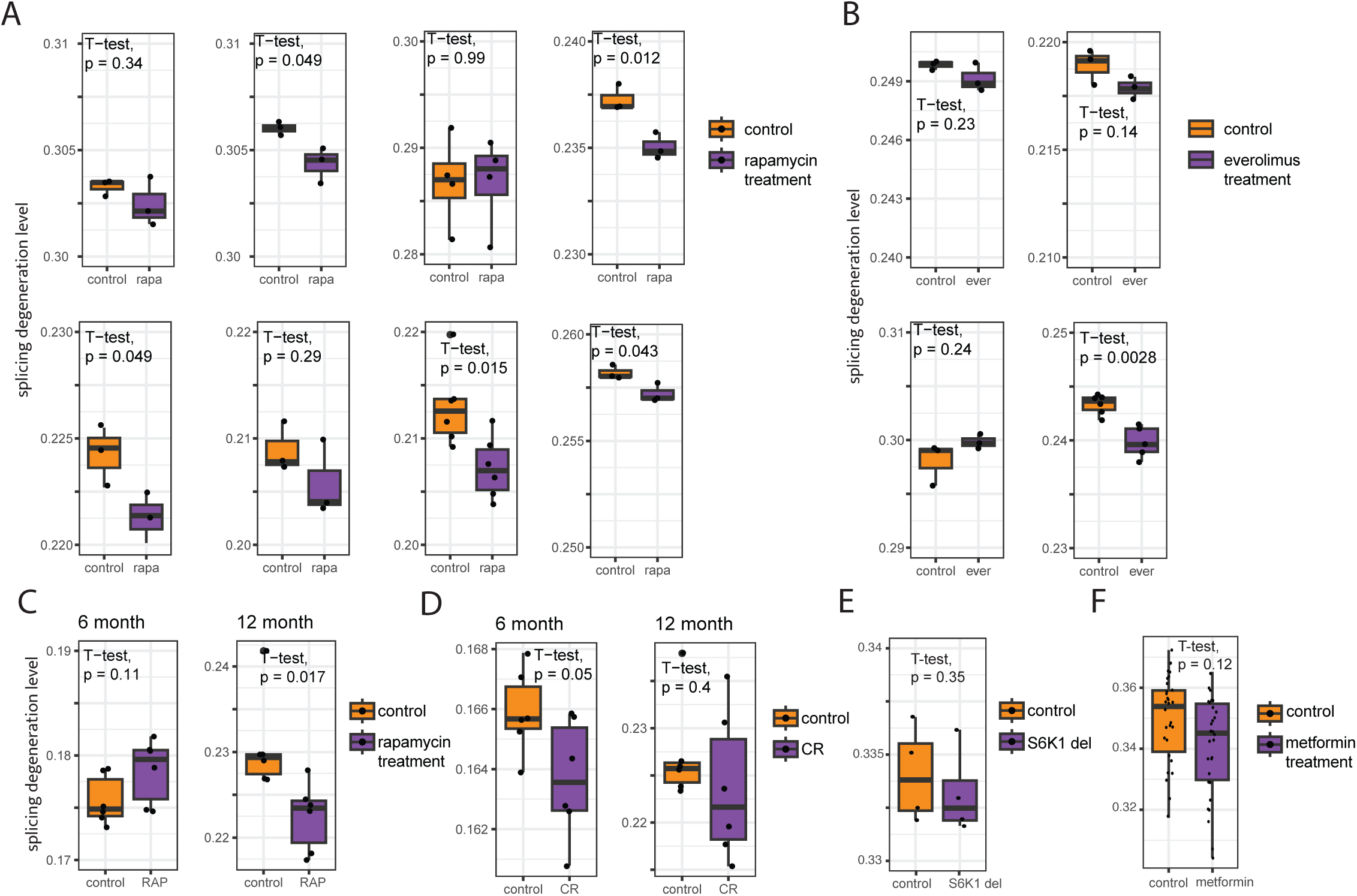
Longevity interventions can restore aging-related splicing degeneration. (A) Splicing degeneration level changes in human cells with and without rapamycin treatment. (B) Splicing degeneration level changes in human cells with and without everolimus treatment. (C) Splicing degeneration level in 6-month and 12-month years old mice with and without rapamycin treatment. (D) Splicing degeneration level in 6-month and 12-month years old mice subjected or not to calorie restriction. (E) Splicing degeneration level in wide type and p70 S6 protein kinase (S6K1) knockout mouse cortex. (F) Splicing degeneration level changes in human cells with and without metformin treatment.

Calorie restriction is another established longevity intervention ^38–40^. We examined the effects of calorie restriction on mouse data; interestingly, the results showed a significant reduction in splicing degeneration levels in younger mice and a reduction, albeit not significant, in aged mice (Figure 4D). Several previous studies highlighted the potential of metformin in retarding aging and alleviating aging-related diseases by targeting molecules implicated in the aging process ^41,42^. We also examined the splicing degeneration level in cells subjected to metformin or S6K1 deletion, which also exhibits the features of slowed aging ^43^. The results revealed a possible decrease in splicing degeneration levels following the treatment, although the difference was not significant (Figure 4E-F). Collectively, all of the above analyses support the idea that the aging-related splicing degeneration may be reversed following treatment with established longevity interventions. Thus, the splicing degeneration level may serve as a reliable biomarker of aging from the perspective of damage accumulation, showing the potential to uncover new aging-related processes and events.

### Identification of SFs that regulate splicing degeneration in aging

In order to explore mechanisms underlying splicing degeneration during the aging process, we then focused on splicing factors (SFs), as these proteins play a pivotal role in regulating AS, e.g. by binding to pre-mRNA and acting as splicing enhancers or silencers, thereby regulating alternative splicing ^14^. Using a linear regression model similar to the identification of aging-related AS events with the data from GTEx, we revealed the SFs exhibiting altered gene expression levels with age (Figure S10A, Supplementary Table 2). Notably, the expression levels of numerous SFs demonstrated aging-associated changes; however, the observed trends diverged across different tissues, and brain tissues were clustered into distinct categories from other tissues (Figure S10B). These tissue-specific variations in SF expression levels suggest a tissue-specific AS regulatory landscape during aging. Since gene ontology of the genes harboring aging-related AS events (Figure 2G) showed a significant enrichment in the function of RNA splicing, we presumed the splicing changes of SFs may also play a regulatory role in aging. To test this hypothesis, we identified the aging-related AS events that occur at SF genes and defined the SFs exhibiting altered expression levels with advancing age as gene-SFs, and SFs exhibiting altered splicing patterns with advancing age as AS-SFs (Figure S11A). Among the top 100 gene-SFs and top 100 AS-SFs, 29 SFs were found to be shared (Figure 5A, Supplementary Table 3), for example, PUF60, whose gene expression level and splicing level (exon skipping of the fifth exon) correlated with age (Figure S11A). To explore the regulation effect of SFs on aging- related AS events, we calculated the correlation between PSI values of aging-related AS events and the gene expression levels (TPMs) of each gene-SF or PSI values of each AS-SF. The results showed that there is a stronger correlation between the PSI values of SFs and aging-associated AS events, which indicates that the AS of SFs also has an important regulation effect on aging-related splicing changes (Figure 5B). There is an apparent “self-regulation” of SFs, wherein the AS change of SFs itself contributes to the overall AS change during the aging process. Hence, in the following analyses, we also included the SFs with aging-related AS changes. Through the protein-protein interaction network of the SFs whose gene expression or splicing changes with age, several key factors emerged, including SR proteins–SRSF1, SRSF3, and heterogeneous ribonucleoproteins (hnRNPs)–HNRNPA1, HNRNPH1 (Figure 5C).

**Figure 5.**
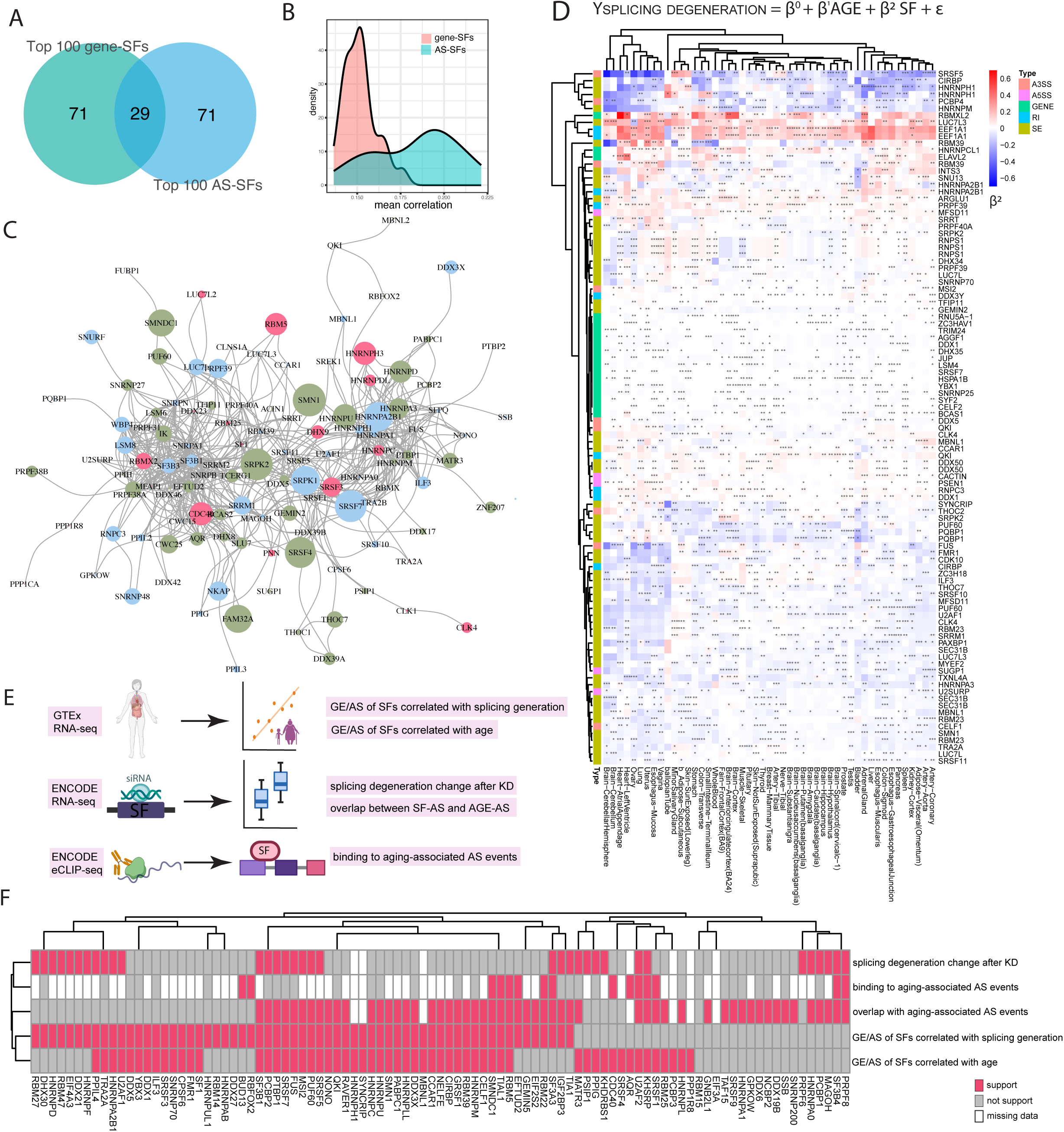
Identification of splicing factors that regulate splicing degeneration in aging. (A) Overlap between the top 100 SFs whose gene expression levels are correlated with age (gene-SFs) and the top 100 SFs whose AS correlates with age (AS-SFs) (data from GTEx). (B) Distribution of the average of the Pearson correlation coefficient between the AS level of aging-related AS events and the gene expression level of SFs (pink) or the AS level of SFs (green). (C) Protein-protein interaction network of the union of the SFs in Figure 5A. (D) Upper: Linear regression model for quantifying the contribution of SFs to splicing degeneration. ‘SF’ indicates the TPM of SF expression level or PSI values of SF AS level. Lower: Heatmap of the coefficient of SF in the linear regression model among different tissues. The colors indicate the coefficient of ‘SF’ in the linear regression model, with darker colors indicating higher coefficients. Blue: the gene expression or splicing level of specific SF is negatively correlate with splicing degeneration level; Red: the gene expression or splicing level of specific SF is positively correlate with splicing degeneration level. (E) Workflow to study regulatory roles of SFs on aging-related splicing degeneration. (F) Heatmap shows the relationship between each splicing factor and evidence for the regulatory role of splicing dysregulation in aging. Each pink square indicates that a SF has a regulatory role in the analysis of that type of evidence. Gray squares indicate the opposite. White squares show that this SF could not be detected due to the lack of or insufficient sequencing data.

We further focused on the relationship between SFs and splicing degeneration level and found that numerous SFs have a significant correlation with the splicing degeneration level (Figure S11B-C). Due to the splicing degeneration level and the expression level / AS level of specific SFs both changing with age, to understand the regulatory role of splicing factors on splicing degeneration and eliminate the effects of age, we used a linear regression model to counteract the effects of aging and identified key SFs contributing to the regulation of splicing degeneration (Figure 5D, and Methods). After eliminating the effect of age, there were still many SFs that showed a significant regulatory effect on the splicing degeneration level (Figure 5D). Since the brain is the least affected by splicing degeneration with age (Figure 3B) and exhibits the strongest negative correlation between SF expression and age (Figure S10A), we further investigated the impact of SFs on brain splicing. In Figure 5D, three genes (ELAVL2, HNRNPCL, and RBMXL2) show a positive correlation with splicing degeneration levels, with these genes being particularly noteworthy. We examined the expression levels of these three SFs in brain tissues and found that two were significantly downregulated during aging, while their expression remained unchanged in other tissues (Figure S12A–F). Expanding this analysis to all SFs, we identified several that positively regulate splicing degeneration and are consistently downregulated with age. Notably, multiple SFs exhibit this pattern (Figure S12G), suggesting a potential mechanism underlying the stable splicing degeneration levels observed in the brain.

To further understand the functional role of these SFs on splicing degeneration, we analyzed the existing large-scale RNA-seq datasets from the ENCODE consortium with knockdown of 96 splicing factors. For each SF, we identified AS events that are significantly altered upon SF knockdown and checked the overlap with aging-related AS events (Figure S13A). We also calculated the splicing degeneration level changes in SFs knockdown samples and control samples, to get the SFs with the regulatory effect on aging-related splicing degeneration (Figure S13B). Moreover, in order to capture SFs that directly act on aging-related splicing abnormalities, for the candidate SFs, we identified their binding sites within the region of aging-related AS events using eCLIP-seq data from ENCODE (Figure S13C). By combining all evidence from RNA interference sequencing (RNAi-seq) and eCLIP-seq, and the above results of the SF changes during aging and also the analyses of SFs on regulating splicing degeneration level (generated using GTEx data), we comprehensively demonstrate the role of SFs in regulating aging-related splicing dysregulation (Figure 5E-F).

Many identified SFs highlight the important regulatory role of AS in aging. For example, the poly(U) binding splicing factor 6 (*PUF60*), whose expression level is negatively correlated with age in most tissues (41 out of 54 tissues), also exhibits a negative correlation with splicing degeneration level (Figure S10D). The skipping levels of *PUF60* exon 2 and exon 5 also correlated with overall splicing degeneration levels. Moreover, there is a significant overlap between *PUF60*-regulated AS events and aging-related AS events, and the splicing degeneration level was reversed after the knockdown of *PUF60* (Figure S13D). All of the above evidence indicates that the low expression of *PUF60* contributes to splicing degeneration in aging.

In summary, we have identified SFs that play important regulatory roles in aging-related splicing disorders and uncovered a “self-regulation” model of several SFs. The comprehensive understanding of aberrant splicing mechanisms during the aging process contributes to a deeper understanding of the fundamentals of aging and facilitates the development of future longevity interventions.

### Abnormalities in SFs influence age-related transcriptome changes

In order to further understand the regulatory role of SFs in the aging process, we employed recently established multi-tissue transcriptomic clocks of chronological age, lifespan-adjusted age and mortality trained on over 6,000 gene expression samples from mouse, rat and human tissues. Applying the clocks to each SF knockdown sample, we compared their transcriptomic age (tAge) with the control samples ^44^. The results showed that out of 88 SFs examined, 68 showed an elevation in tAge after SF knockdown, with 36 knockdowns exhibiting a statistically significant increase of tAge (BH-adjusted p-value, quantified by lifespan-adjusted clock < 0.05; Figure 6A). The increased transcriptome age caused by SFs knockdown illustrates the important role of SF in the aging process and lifespan. As a control, we also quantified the tAge changes induced by the knockdown of other RNA binding proteins (RBPs) and compared the distribution of tAge changes among RBPs, SFs and the key SFs that regulated splicing degeneration (SFs exhibiting at least three positive pieces of evidence across the five aspects are shown in Figure 5F). Notably, our results demonstrate a significantly larger transcriptomic age alteration subsequent to SF knockdown compared to other RBPs, with the key SFs implicated in splicing degeneration having the highest effect (Figure 6B). Similar results were also observed according to chronological and mortality clocks (Figure S14). This evidence underscores the consequential implications of SF dysregulation on the aging process and the attendant diminution of health span.

**Figure 6.**
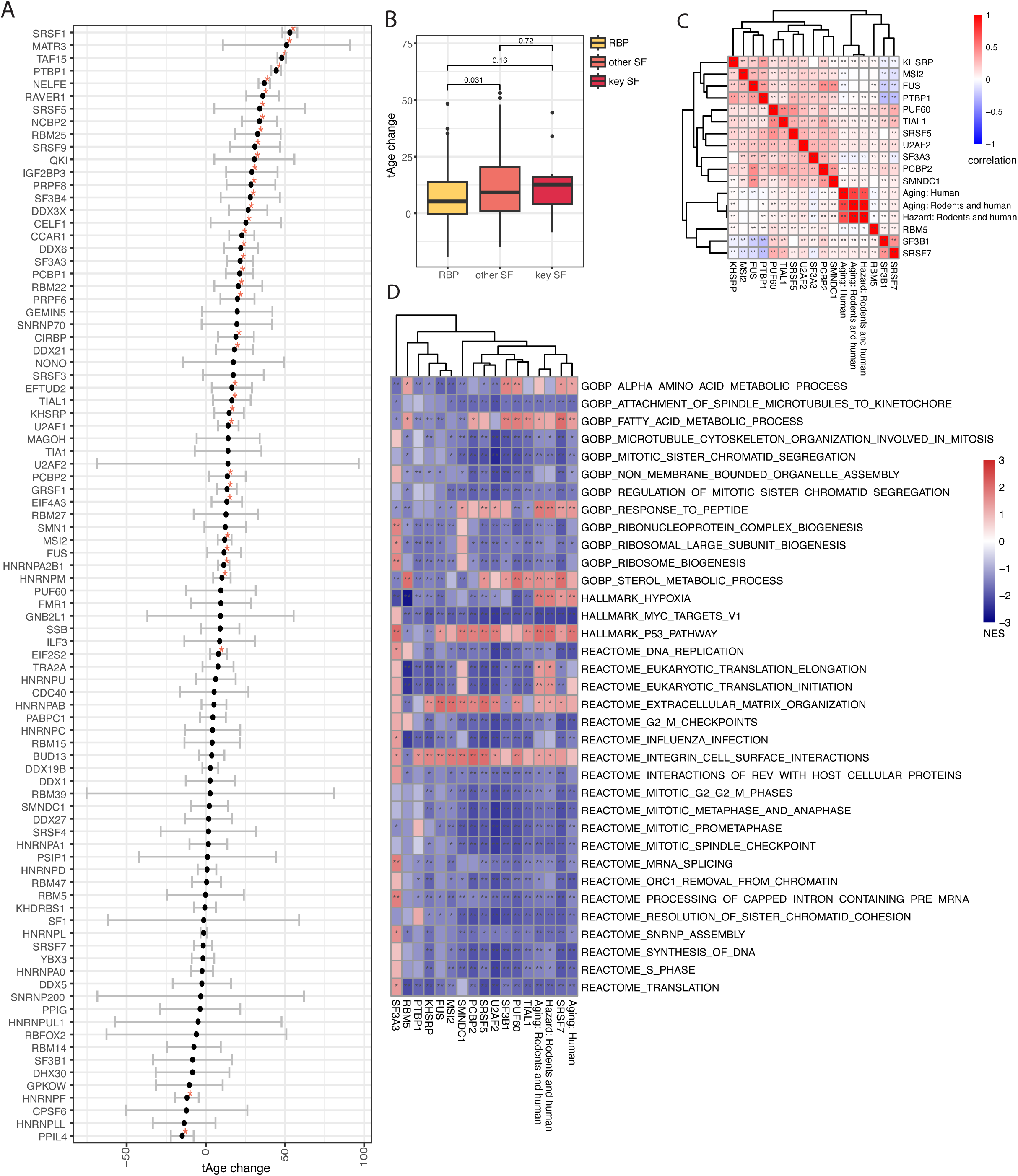
Relationship between SFs and aging-related transcriptome signatures. (A) Transcriptome age (lifespan-adjust clock) changes following each SF knockdown relative to control. (*: P<0.05, Student’s t-test) (B) Distribution of transcriptome age (lifespan-adjust clock) change following the conditions of the knockdown of key SFs that regulated splicing degeneration, other SFs, and other RBPs. (C) Heatmap of the Pearson correlation coefficients between gene expression changes induced by key SF knockdown and signatures of aging. The color intensity and gradient denote the strength and direction of the correlation, with darker colors indicating stronger correlations. Positive correlations are shown in red, while negative correlations are shown in blue. (*: P<0.05, **: P<0.01, Pearson’s correlation test). The SFs that have at least four positive pieces of evidence over the five aspects of evidence in Figure 5 are shown. (D) Functional GSEA pathways enriched by gene expression changes induced by key SFs knockdown and aging (*: P<0.05, **: P<0.01). The SFs that have at least four positive pieces of evidence over the five aspects of evidence in Figure 5 are shown.

Among the SFs, SRSF1, a member of the serine and arginine-rich (SR) protein family, emerged as exerting the most pronounced influence on the transcriptomic age in response to KD. This observation nicely aligns with recent findings, that overexpression of SRSF1 could reprogram cells to a more youthful state ^30^. This also verifies the utility of transcriptome age as a metric for investigating the interplay between SFs and aging. Several other SFs, such as SRSF5, PTBP1, QKI, SRSF5, and U2AF1, which exhibited substantial alterations in response to knockdown, also have an important regulatory role in aging and lifespan.

To further study the critical role of SFs on aging, we examined the relationship between SF-related gene expression changes and multi-tissue aging and mortality signatures of humans and rodents ^6^. The majority of SFs demonstrated significant positive correlations with aging signatures (Figure S15); several key SFs that regulated splicing degeneration are shown in Figure 6C. Gene set enrichment analyses (GSEA) showed that these SFs may operate through MYC and P53 pathways, and ribosome-related translation (Figure 6D). In summary, our findings reveal a close relationship between SFs and aging, particularly those SFs that regulate splicing degeneration.

## Discussion

Aging is a complex biological process characterized by numerous changes at the molecular, cellular, organ, and organismal levels ^1,45^. As we strive to extend human lifespan and cure age-related diseases, understanding the nature of aging and mechanisms underlying this process is crucial. Various aging hallmarks have been previously defined, and our study suggests that AS emerges as a novel player in this intricate landscape.

AS is an important biological process during gene expression, allowing a single gene to produce multiple protein isoforms ^8^. Dysregulation of AS has been implicated in various diseases, especially aging-related diseases ^46,47^, hence the role of AS in aging has gained much attention recently ^48,49^. However, in previous studies, researchers often concentrated solely on identifying aging-associated AS events ^22^, or focused exclusively on individual splicing factors or in single tissues ^25,50^. Here, we explored the overarching remodeling of AS during aging, studied how splicing affects protein functions throughout the aging process and revealed its potential as a molecular indicator for longevity interventions. Through comprehensive analyses of thousands of RNA-seq data from both human and mice, we found that genome-wide splicing degeneration critically contributes to aging. This degeneration of AS leads to functional decline and can serve as a new hallmark of aging.

We present a method to quantify splicing degeneration levels, which measures the extent of functional decline caused by dysregulated splicing. Notably, there is a significant positive correlation between splicing degeneration levels and age, whereas the splicing degeneration level can be partially reversed by longevity interventions. There are also certain limitations. First, the AS quantification method is influenced by the sequencing depth and read length, making it difficult to do cross-dataset comparisons, especially when dealing with different sequencing protocols. Moreover, we only focus on the annotated AS events in our results. While prioritizing annotated AS events ensures more robust results, it may lead to the loss of some information related to splicing deterioration. Additionally, the sensitivity of splicing degeneration level quantification requires improvement, as accurately capturing subtle aging-related changes remains challenging.

We demonstrated extensive splicing alterations during aging and identified several potential mechanisms underlying these changes. In particular, we observed that certain SFs play a regulatory role in this process. Additionally, age-related shifts in cell composition and immune cell infiltration may contribute to splicing degeneration. It is important to note that not all splicing alterations arise from splicing errors; some may represent adaptive responses to stress or reflect changes in tissue composition.

Our research established the intricate interplay between aging pathways and AS and extended it to longevity interventions. We specifically focused on rapamycin. This is a well-known longevity-promoting compound that exerts its effect on aging by targeting the MTOR pathway ^51^. Interestingly, this same pathway is closely linked with AS, emphasizing the interconnectedness of these fundamental cellular processes. In our study, we found that numerous misregulated aging-related AS events are restored following rapamycin treatment, and that the overall splicing degeneration level is decreased following rapamycin treatment and other longevity interventions.

To better understand splicing degeneration during aging, we examined the associated mechanisms. We describe a “self-regulation” model of key splicing factors, wherein the AS of the splicing factor itself can regulate the overall splicing degeneration level. We performed a comprehensive analysis of the role of SFs in aging, identifying SFs regulating aging-associated splicing degeneration. We systematically examined the transcriptomic changes following the knockdown of SFs and explored the relationship between SFs and aging signatures. Although we are not surprised that the knockdown of SFs increases the transcriptome age, our analysis clearly pointed out the effect of SF knockdown on predicted age and ranked the effects of SFs, providing information on which SF may be important during aging. Our study contributes to a deeper understanding of the role of SFs in aging, which could also provide insights into other studies in the field.

There are several known aging hallmarks, such as epigenetic changes, genome instability, and mitochondrial dysfunction ^3^. Recognizing AS as a potential biomarker of aging allows to better characterize the aging process. Coupled with other markers, splicing signatures may provide a more complete understanding of aging dynamics and facilitate targeted interventions. Moreover, our findings suggest that splicing restoration is linked to lifespan extension. As we continue to explore the molecular underpinnings of aging, identifying splicing events as therapeutic targets may become increasingly critical. Since AS may be regulated, can AS modulation help combat age-related decline? Our study suggests it’s a promising avenue for future studies.

In summary, our study sheds light on a novel pattern of splicing degeneration during aging, and reveals the intricate relationship between aging and alternative splicing. As we strive to extend lifespan, recognizing the critical importance of elucidating these mechanisms becomes paramount. Our work not only enhances awareness but also paves the way for new longevity interventions in the quest for healthy aging.

## Methods

### AS level quantification and AS clock building

In processing the data retrieved from the GTEx database, the BAM files are used to quantify AS levels through the Paean tool, a parallel computing system operating on GPU-CPU platforms ^28^. The output of Paean is highly consistent with results from traditional AS quantification tools as well as experimental results, which supports its application in our analysis. The Percent Spliced In (PSI) value, which denotes the fraction of mRNAs representing the inclusion isoform, was calculated for each AS event in each sample.

To verify the significant role of AS in aging, we constructed AS aging clocks. For each tissue in GTEx, samples were randomly divided into training and test sets, with 90% allocated to the training set and 10% to the test set. For the training sets, we used PSI values of all AS events as the input for a random forest model and predicted the age in the test set. This procedure was repeated 100 times, and the mean correlations between predicted and true ages were plotted.

### Differential AS analysis in aging samples

We employed several approaches to identify differential AS events between young and old samples. Using the splicing patterns of young samples as a baseline, we compared the PSI values of AS events in aged samples with the average PSI levels of young samples. AS events with |ΔPSI| > 0.1 and *P* < 0.05 were considered differential. Additionally, we applied an alternative method using |log₂FoldChange| > 1 and *P* < 0.05 (one-sample *t*-test) to identify differential AS events and calculate their correlation with age (Figure S2B). To further investigate, we used a differential expression approach (via the limma package in R) to compare AS events between each old and young groups, applying criteria of |log₂FoldChange| > 1 and FDR < 0.05.

### Identification of aging-associated AS events in GTEx dataset

In order to identify aging-associated alternative splicing events in each tissue, we set up a linear regression model as follows:

Ypsi = β^0^ + β^1^AGE + β^2^ GENDER + β^3^RACE + β^4^DTHHRDY + ε in which β^0^ is the intercept for the model, β1 is the coefficient of age, β2 is the coefficient of sex, β2 is the coefficient of race, β3 is the coefficient of race, β4 is the coefficient of type of death classification, and ε is the error.

To enhance the robustness of our findings, we implemented stringent criteria that need to meet the following three conditions: (1) P value for β1 (coefficient of “AGE” in the linear regression model) < 0.05, (2) The P value for the Pearson correlation test between PSI value and age < 0.05, (3) Consistency in the directionality of β1 and the Pearson correlation coefficient, with both being either more than 0 or less than 0. Through the application of these cutoffs, we successfully identified numerous AS events in each tissue. The aging-associated AS events across tissues are defined as the AS events associated with age in at least 10 tissues.

### Generalized Summary-data-based Mendelian Randomization Analysis

GSMR is a powerful tool for inferring causal associations between exposures and outcomes using summary-level data from genome-wide association studies (GWAS) ^29^. We obtained summary statistics of PSI values for aging-associated AS events from the Genotype-Tissue Expression (GTEx) project ^52^, which were used as instrumental variables (IVs) in the GSMR analysis. We included all tissues that exhibit at least one AS event for a given gene (based on FDR). We selected two aging-related outcomes: the Horvath epigenetic age ^53^ and parental lifespan ^54^. The summary statistics for these outcomes were obtained from publicly available GWAS data.

To ensure the validity of the IVs, we applied several filtering steps. First, we selected only bi- allelic SNPs with a minor allele frequency (MAF) ≥ 0.01 and an imputation quality score (INFO) ≥ 0.9. Second, we removed SNPs in linkage disequilibrium (LD) with an r^2^ threshold of 0.01 using the clumping function in PLINK (v1.90) ^55^. Third, we excluded SNPs associated with the outcomes at a genome-wide significance level (P < 5 × 10⁻⁸) to avoid horizontal pleiotropy. We performed the GSMR analysis using the GCTA software (v1.93.1) ^56^ with default parameters. The causal effects of AS events on the Horvath Age and parental lifespan were estimated, and the significance threshold was set at a Bonferroni-corrected P-value < 0.05, accounting for the number of tested AS events.

### Feature analyses of aging-associated AS events

To examine conservation of alternative regions of aging-associated AS events, we used human phastCons data from alignments of 100 placental mammals, which was downloaded from UCSC (http://genome.ucsc.edu/). For each AS event, the average phastCons score of each site was calculated. The splice site scores were calculated using the MaxEntScan (http://hollywood.mit.edu/burgelab/maxent/Xmaxentscan_scoreseq.html). Specifically, we use sequences at positions (-3 to +6, last 3 bases of exons and first 6 bases of succeeding intron) of the 5’ ss, which have the GT sequences at the positions +1, +2, and the sequences at positions - 20 to +3 of the 3’ss with the AG consensus at position -2, -1. The sequences are analyzed with MaxEntScan that applies the Maximum Entropy model to calculate splice site strength ^57^. The GO analysis for the human genes harboring aging-related AS events was built using R package “clusterProfiler” ^58^.

### Classification of AS events into “damaging” and “non-damaging” AS events

In order to study functional consequences of aging-associated AS events, we annotated each AS event into “damaging” and “non-damaging”. For SE, A3SS and A5SS, we examined whether the length of the alternative fragment is multiple of three. AS events whose alternative fragment length is not a multiple of three can cause frameshift. For the AS events whose alternative fragment length is a multiple of three, we examined whether the alternative region overlaps with a protein domain region (annotation for protein domain was downloaded from Uniprot). The AS events which can cause frameshift or locate at a domain region were classified to “damaging” consequences. For IR events, we identified the stop codon in the specific intron and checked if there is at least one stop codon within the reading frame. The IR events which can cause a PTC are classified to “damaging” AS events (Figure 2E).

### Quantification of splicing degeneration level

To quantify the overall splicing degeneration level from PSI values of AS events, we only used the damaging AS events. Specifically, we identified three categories indicative of AS events with damaging sequences: (1) A premature stop codon; (2) Frameshift; and (3) Loss of a peptide sequence within a protein domain region. Employing the cutoff with a PSI value > 0.3 for damaging IR events and PSI values < 0.7 for damaging SE, A3SS, and A5SS events, we computed the proportion of these types of AS events as the measure of splicing degeneration level (Figure 3A).

### RNA-sequencing analyses of mouse datasets

For the mouse datasets used in this research, we downloaded raw data from GEO (with accession numbers: GSE92486 ^59^, GSE127758, GSE155407 ^60^, GSE121395 ^61^, GSE83931 ^62^, GSE175633^63^, GSE145480 ^64^, GSE175854 ^65^. RNA-seq reads were aligned to the mouse genome (mm10) using STAR with the parameters “--chimSegmentMin 2 --outFilterMismatchNmax 3 --alignEndsType EndToEnd”, and PSI values were calculated using rMATS ^66^. To identify the aging-associated AS events, a cutoff of |ΔPSIavg | > 0.1 and P < 0.05 were used. For the quantification of splicing degeneration levels, a similar method to the one described above was used.

### Longevity intervention analyses

For the longevity intervention data generated from human cells or mouse, raw RNA sequencing data were downloaded from GEO (accession numbers: GSE107965 ^67^, GSE184427 ^68^, GSE225751 ^69^, GSE89774 ^70^, GSE137353 ^71^, GSE225683 ^72^, GSE225751 ^69^, GSE123740 ^73^, GSE99875, GSE143659, GSE90755^74^, GSE131901), the RNA sequencing reads were aligned to the human genome (hg38) or mouse genome (mm10) using STAR with specified parameters “--chimSegmentMin 2 --outFilterMismatchNmax 3 --alignEndsType EndToEnd”, and PSI values were calculated using rMATS ^66^. To identify AS changes following longevity interventions, we conducted comparisons between the treatment group and its paired control samples and selected the AS events exhibiting significant differences (with a cutoff of |ΔPSIavg| > 0.1 and FDR < 0.05) as the AS events related to the intervention. For the identification of rapamycin-associated AS events, we defined it as the AS events associated with rapamycin in at least three datasets.

### RNAi-seq and eCLIP seq data analyses

RNA-seq data from the knockdown of 227 RBPs and the eCLIP sequencing data were downloaded from ENCODE consortium. For RNAi-seq, the sequencing reads were aligned to the human genome(hg38) using STAR and PSI values were calculated using rMATS ^66^. Gene expression levels of each sample were quantified by Expectation Maximization (RSEM) ^75^ and differential gene analysis was generated by R package ‘DESeq2’. For the compilation of SFs, we adopted a previous study which combined three databases of SFs and selected the most reliable as the final SF list ^76^. For each SF, we compared the changes between SF knockdown and paired control samples to identify the AS events with significant changes (with cutoff of |ΔPSIavg| > 0.1 and P < 0.05), which were defined as SF-associated AS events. We also calculated the splicing degeneration level of each sample and compared between splicing factor knockdown and control groups. For eCLIP sequencing, we downloaded the bed files containing the peaks generated by CLIPper ^77,78^. We then mapped the binding site of each SF to aging-associated AS events to study the regulation role of SFs.

### Analysis of regulatory roles of SFs on aging-related AS change

For the identification of top 100 gene-SFs and splicing-SFs in Figure 5A, we first calculated the correlation between age and SF gene expression or AS level of SFs in each tissue separately. We then ranked the mean of the absolute correlation values of correlations in all tissues and selected the top 100 (Figure S11A).

In order to comprehensively demonstrate the role of SFs in regulating aging-related splicing dysregulation, we conducted the analyses from five different aspects (Figure 5E). (1) Using data from GTEx, we examined whether the gene expression or AS of a specific splicing factor is associated with age in at least 15 tissues. (2) Using data from GTEx, we examined whether the gene expression level or AS of a specific splicing factor correlated with the splicing degeneration level in at least 15 tissues. (3) Using RNA sequencing data from ENCODE, we examined whether the splicing degeneration level is significantly changed between SF knockdown samples and control samples. (4) Using RNA sequencing data from ENCODE, we examined whether there is a significant overlap between SF-associated AS events and aging-associated events. (5) Using eCLIP sequencing data from ENCODE, we examined whether the specific splicing factor binds to at least 10% aging-associated AS events. Combining all these five aspects, we generated a figure to show the results, wherein a pink square indicates that a splicing factor passes the test and gray shows it does not pass (Figure 5F). The SFs showed at least three positive over the five aspects of evidence are defined as key SFs that regulated splicing degeneration, including PPIL4, TRA2A, HNRNPA2B1, U2AF1, SF3B1, PCBP2, PTBP1, SRSF7, FUS, MSI2, PUF60, SRSF5, NONO, QKI, RAVERA, HNRNPC, HNRNPU, SMN1, PABPC1, HNRNPLL, DDX3X, CCAR1, MELFE, CIRBP, GRSF1, RBM39, HNRNPM, CELF1, SMNDC1, TIAL1, RBM5, EFTUD2, RBM22, SF3A3, IGF2BP3, TIA1, MATR3, U2AF2, KHSRP, SRSF1, SF3B4 and PRPF8.

### Transcriptomic clock and aging signature analyses of SFs

To assess the transcriptomic age (tAge) changes induced by SFs knockdowns, we utilized multi-species (trained on mice, rodents, and humans) multi-tissue transcriptomic clocks of (1) chronological age, (2) lifespan-adjusted clock (i.e. chronological age divided by expected maximum lifespan), and (3) expected mortality (according to chronological age and corresponding Gompertz mortality curve). While chronological clock was trained only on control healthy samples, lifespan-adjusted and mortality clocks were developed based on healthy mice, rats and humans, along with animals subjected to various lifespan-shortening and longevity interventions ^44^.

Filtered RNA-seq data representing SD knockdowns and corresponding controls was log-transformed and scaled, and control samples were used as a reference group to calculate relative gene expression profile for every sample. The missing values corresponding to clock genes not detected in the data were imputed with the precalculated average values. To assess correlation of logFC induced by SD knockdowns (compared to control samples) and gene expression signatures of aging and hazard established in our previous studies^6,79^. The utilized signatures included (1) multi-tissue signature of chronological age in humans, (2) in humans and rodents, and (3) multi-tissue signature of mortality in humans, mice and rats. Pairwise Spearman correlation for gene expression changes induced by SF knockdown as well as transcriptomic signatures of aging/lifespan was calculated based on the union of top 650 genes with the lowest p-value for each pair of signatures.

For the identification of enriched functions affected by SF knockdown, we performed functional GSEA ^80^ on a pre-ranked list of genes. Gene ontology: biological process (GObp), HALLMARK, KEGG and REACTOME ontologies from the Molecular Signature Database (MSigDB) were used as gene sets for GSEA. The GSEA algorithm was performed separately for each SF knockdown and age group via the fgsea package in R with 5000 permutations.

## Acknowledgments

We thank Dr. Zefeng Wang (Southern University of Science and Technology) and members of the Gladyshev lab for discussion. This work was funded by National Institute on Aging and Impetus grants.

## Supplementary Figure legends

**Figure S1.**

(A) Correlation between age and the number of different types of differential AS events in aging samples in each human tissue. Data from GTEx. A3SS: alternative 3’ splice site, A5SS: alternative 5’ splice site, RI: retain intron, SE: skip exon. (B) Correlation between age and the number of differential AS events in indicated aging-related pathways. Data from GTEx. Asterisks indicate significance of the correlation. *: P<0.05, **: P<0.01, ***: P<0.001.

**Figure S2.**

(A) Correlation between the number of differential AS events and age in the indicated human tissues from GTEx. PSI values for each AS event identified in each aged sample are compared with the average PSI levels of young samples using the different cutoffs. From left to right: Old group: older than 40 years, young group: younger than 35 years; Old group: older than 35 years, young group: younger than 35 years; Old group: older than 40 years, young group: younger than 40 years; Old group: older than 35 years, young group: younger than 30 years; For each old sample, AS events with |delta PSI value| > 0.1 and P < 0.05 were identified as differential AS events. Asterisks indicate the significance of the Pearson correlation coefficient, *: P<0.05, **: P<0.01, ***: P<0.001. (B) Correlation between the number of differential AS events and age in indicated human tissues from GTEx. Asterisks indicate the significance of the Pearson correlation coefficient (*: P<0.05, **: P<0.01, ***: P<0.001). (C) Counts of differential AS events in each age groups compared with young group.

**Figure S3.**

(A) Tissue-specific alternative splicing clocks which were established using PSI values of all AS events. In each tissue, 90% of the samples were used to train a random forest model and 10% of the samples were used to test the model. (B) Scatter plots of the correlation between chronological age and predicted age in selected tissues. Training and test sets are denoted by color. Pearson correlation coefficient and P value are shown in the text.

**Figure S4.**

Counts of different types of aging-associated AS events in each tissue. Different colors indicate positive or negative correlations with age. A3SS: alternative 3’ splice site, A5SS: alternative 5’ splice site, RI: retain intron, SE: skip exon.

**Figure S5.**

(A) Distribution of P-values for the causal effects of sQTLs on Horvath Age or lifespan in each tissue, as estimated by GSMR. Genes exhibiting significant causal effects in at least 15 tissues are shown. Point colors represent different tissues. (B) Scatter plot of significant causal effects of sQTLs on Horvath Age and lifespan in each tissue, as generated by GSMR.

**Figure S6.**

(A)The P values distribution of the causal effect (generated by GSMR) of aging-related AS on Horvath Age or lifespan in each tissue. The genes showing significant causal effects in at least 15 tissues are shown. The color of points represent different tissues. (B) Scatterplot of significant causal effect (generated by GSMR) of aging-related AS on Horvath Age and lifespan in each tissue. (C-D) The heatmap of P values of the causal effect (generated by GSMR) of aging-related AS on Horvath Age (C) or lifespan (D) in each tissue. The color denotes the - log P values. The genes showing significant causal effects in at least 5 tissues are shown. * :P < 0.05.

**Figure S7.**

Gene ontology of genes harboring aging-associated AS events in each tissue.

**Figure S8.**

(A) Trend of mean PSI values (PSI values in SE, A3SS, A5SS and 1-PSI value in RI) of all AS changes during aging in all tissues (left) and each different tissue (right). (B) Correlation between splicing degeneration level and age in each tissue under the different use of cutoffs when quantifying the splicing degeneration level. (C) Correlation between splicing degeneration level (only use PTC and frameshift types) and age in each human tissue (data from GTEx). Asterisks indicate significance of the Pearson correlation, *: P<0.05, **: P<0.01, ***: P<0.001. (D) Splicing degeneration level in tumor and adjacent normal samples (data from TCGA). We only use the cancer types with at least 20 normal samples. BRCA: Bladder Urothelial Carcinoma, COAD: Colon adenocarcinoma, HNSC: Head and Neck squamous cell carcinoma, KICH: Kidney Chromophobe, KIRC: Kidney renal clear cell carcinoma, KIRP: Kidney renal papillary cell carcinoma, LIHC: Liver hepatocellular carcinoma, LUAD: Lung adenocarcinoma, LUSC: Lung squamous cell carcinoma, PRAD: Prostate adenocarcinoma, STAD: Stomach adenocarcinoma, THCA: Thyroid carcinoma, UCEC: Uterine Corpus Endometrial Carcinoma.

**Figure S9.**

(A) Overlap between aging-related AS events and rapamycin-associated AS events. Fisher’s exact test was used to calculate the significance of the overlap. (B) Gene ontology of genes harboring rapamycin-associated AS events. (BP: Biological Process, CC: Cellular Component, MF: Molecular Function). (C) Density of PSI values distributions of each type of rapamycin-associated AS events. Different colors indicate different datasets.

**Figure S10.**

(A) Counts of splicing factors whose gene expression level correlated with age in each tissue. Different colors indicate positive or negative correlation with age. (B) Heatmap of the correlation between gene expression level of all SFs and age in each tissue.

**Figure S11.**

(A) Workflow for identifying top 100 gene-SF and splicing-SF in Figure 5A (top) and the correlation between age and PUF60 expression level and splicing level as an example (bottom). (B) Heatmap of the correlation between TPM of splicing factors and PSI values of aging-associated AS events in each tissue. (C) Heatmap of the correlation between PSI values of all AS events in SFs and PSI values of aging-related AS events in each tissue.

**Figure S12.**

(A) Correlation between ELVAL2 expression levels and age in brain tissues.(B) Correlation between HNRNPCL expression levels and age in brain tissues. (C) Correlation between RBMXL2 expression levels and age in brain tissues. (D) Correlation betweenELVAL2 expression levels and age in other tissues. (E) Correlation between HNRNPCL expression levels and age in other tissues. (F) Correlation between RBMXL2 expression level and age in other tissues. (F) The SFs which show positive effect on splicing degeneration level and negative correlation with age in brain tissues. The x-axis represents the coefficient for splicing degeneration level (adjusted for age effects).

**Figure S13.**

(A) Significance of the overlap between aging-associated AS events and the AS events changed after knockdown of each SF (Fisher’s exact test). (B) Significance of splicing degeneration level change after SF knockdown (Student’s t-test). (C) Number of aging-related AS events with the binding of each SF. (D) Correlation between age and gene expression of PUF60 in each tissue (left), correlation between splicing degeneration level and gene expression of PUF60 in each tissue (median) and the splicing degeneration change in PUF60 knockdown cells compared with control samples (right).

**Figure S14.**

(A) Transcriptome age changes (chronological age clock) following the knockdown of each SF. (B) Transcriptome age change (mortality age clock) following the knockdown of each SF. (C) Distribution of transcriptome age changes following the key SFs knockdown, SFs knockdown and other RBPs knockdown. Top panel: chronological age clock; Bottom panel: hazard age clock. (*: P<0.05, Student’s t-test).

**Figure S15.**

Heatmap of the Pearson correlation coefficients between gene expression changes induced by SF knockdown and signatures of aging. The color intensity and gradient denote the strength and direction of the correlation, with darker colors indicating stronger correlations. Positive correlations are shown in red, while negative correlations are shown in blue.

**Supplementary Table 1.**

Aging-related AS events across human tissue.

**Supplementary Table 2.**

The splicing factors exhibited altered gene expression levels with age.

**Supplementary Table 3.**

The list of the top 100 SFs whose gene expression changes with age and the top 100 SFs whose splicing changes with age.

**Supplementary Table 4.**

The corresponding AS events of the gene name in Figure 5D.

## References

1. Gladyshev, V.N., Kritchevsky, S.B., Clarke, S.G., Cuervo, A.M., Fiehn, O., de Magalhães, J.P., Mau, T., Maes, M., Moritz, R., Niedernhofer, L.J., et al. (2021). Molecular Damage in Aging. Nat Aging 1, 1096–1106. 10.1038/s43587-021-00150-3.

2. Gladyshev, V.N. (2016). Aging: progressive decline in fitness due to the rising deleteriome adjusted by genetic, environmental, and stochastic processes. Aging cell 15, 594–602.

3. López-Otín, C., Blasco, M.A., Partridge, L., Serrano, M., and Kroemer, G. (2013). The hallmarks of aging. Cell 153, 1194–1217. 10.1016/j.cell.2013.05.039.

4. de Magalhães, J.P., Curado, J., and Church, G.M. (2009). Meta-analysis of age-related gene expression profiles identifies common signatures of aging. Bioinformatics 25, 875–881. 10.1093/bioinformatics/btp073.

5. Tikhonov, S., Batin, M., Gladyshev, V.N., Dmitriev, S.E., and Tyshkovskiy, A. (2024). AgeMeta: Quantitative Gene Expression Database of Mammalian Aging. Biochemistry (Mosc) 89, 313–321. 10.1134/s000629792402010x.

6. Tyshkovskiy, A., Ma, S., Shindyapina, A.V., Tikhonov, S., Lee, S.G., Bozaykut, P., Castro, J.P., Seluanov, A., Schork, N.J., Gorbunova, V., et al. (2023). Distinct longevity mechanisms across and within species and their association with aging. Cell 186, 2929–2949.e2920. 10.1016/j.cell.2023.05.002.

7. Frenk, S., and Houseley, J. (2018). Gene expression hallmarks of cellular ageing. Biogerontology 19, 547–566. 10.1007/s10522-018-9750-z.

8. Wang, E.T., Sandberg, R., Luo, S., Khrebtukova, I., Zhang, L., Mayr, C., Kingsmore, S.F., Schroth, G.P., and Burge, C.B. (2008). Alternative isoform regulation in human tissue transcriptomes. Nature 456, 470–476. 10.1038/nature07509.

9. Pan, Q., Shai, O., Lee, L.J., Frey, B.J., and Blencowe, B.J. (2008). Deep surveying of alternative splicing complexity in the human transcriptome by high-throughput sequencing. Nat Genet 40, 1413–1415. 10.1038/ng.259.

10. Jutzi, D., and Ruepp, M.D. (2022). Alternative Splicing in Human Biology and Disease. Methods Mol Biol 2537, 1–19. 10.1007/978-1-0716-2521-7_1.

11. Angarola, B.L., and Anczuków, O. (2021). Splicing alterations in healthy aging and disease. Wiley Interdiscip Rev RNA 12, e1643. 10.1002/wrna.1643.

12. Zhang, S., Mao, M., Lv, Y., Yang, Y., He, W., Song, Y., Wang, Y., Yang, Y., Al Abo, M., Freedman, J.A., et al. (2022). A widespread length-dependent splicing dysregulation in cancer. Sci Adv 8, eabn9232. 10.1126/sciadv.abn9232.

13. Matera, A.G., and Wang, Z. (2014). A day in the life of the spliceosome. Nat Rev Mol Cell Biol 15, 108–121. 10.1038/nrm3742.

14. Wang, Z., and Burge, C.B. (2008). Splicing regulation: from a parts list of regulatory elements to an integrated splicing code. Rna 14, 802–813. 10.1261/rna.876308.

15. Wang, N., Hu, Y., and Wang, Z. (2023). Regulation of alternative splicing: Functional interplay with epigenetic modifications and its implication to cancer. Wiley Interdiscip Rev RNA, e1815. 10.1002/wrna.1815.

16. Mazin, P.V., Khaitovich, P., Cardoso-Moreira, M., and Kaessmann, H. (2021). Alternative splicing during mammalian organ development. Nat Genet 53, 925–934. 10.1038/s41588-021-00851-w.

17. Baralle, F.E., and Giudice, J. (2017). Alternative splicing as a regulator of development and tissue identity. Nat Rev Mol Cell Biol 18, 437–451. 10.1038/nrm.2017.27.

18. Mazin, P., Xiong, J., Liu, X., Yan, Z., Zhang, X., Li, M., He, L., Somel, M., Yuan, Y., Phoebe Chen, Y.P., et al. (2013). Widespread splicing changes in human brain development and aging. Mol Syst Biol 9, 633. 10.1038/msb.2012.67.

19. Bhadra, M., Howell, P., Dutta, S., Heintz, C., and Mair, W.B. (2020). Alternative splicing in aging and longevity. Hum Genet 139, 357–369. 10.1007/s00439-019-02094-6.

20. Latorre, E., and Harries, L.W. (2017). Splicing regulatory factors, ageing and age-related disease. Ageing Res Rev 36, 165–170. 10.1016/j.arr.2017.04.004.

21. Heintz, C., Doktor, T.K., Lanjuin, A., Escoubas, C., Zhang, Y., Weir, H.J., Dutta, S., Silva-García, C.G., Bruun, G.H., Morantte, I., et al. (2017). Splicing factor 1 modulates dietary restriction and TORC1 pathway longevity in C. elegans. Nature 541, 102–106. 10.1038/nature20789.

22. Wang, K., Wu, D., Zhang, H., Das, A., Basu, M., Malin, J., Cao, K., and Hannenhalli, S. (2018). Comprehensive map of age-associated splicing changes across human tissues and their contributions to age-associated diseases. Sci Rep 8, 10929. 10.1038/s41598-018-29086-2.

23. Marco, M., Csaba, K., Winona, O., Marta, M., and Vadim, N.G. (2022). Deterioration of the human transcriptome with age due to increasing intron retention and spurious splicing. bioRxiv, 2022.2003.2014.484341. 10.1101/2022.03.14.484341.

24. Holly, A.C., Melzer, D., Pilling, L.C., Fellows, A.C., Tanaka, T., Ferrucci, L., and Harries, L.W. (2013). Changes in splicing factor expression are associated with advancing age in man. Mech Ageing Dev 134, 356–366. 10.1016/j.mad.2013.05.006.

25. Blanco, F.J., and Bernabéu, C. (2012). The Splicing Factor SRSF1 as a Marker for Endothelial Senescence. Front Physiol 3, 54. 10.3389/fphys.2012.00054.

26. Lee, B.P., Pilling, L.C., Emond, F., Flurkey, K., Harrison, D.E., Yuan, R., Peters, L.L., Kuchel, G.A., Ferrucci, L., Melzer, D., and Harries, L.W. (2016). Changes in the expression of splicing factor transcripts and variations in alternative splicing are associated with lifespan in mice and humans. Aging Cell 15, 903–913. 10.1111/acel.12499.

27. Salapa, H.E., Thibault, P.A., Libner, C.D., Ding, Y., Clarke, J.-P.W.E., Denomy, C., Hutchinson, C., Abidullah, H.M., Austin Hammond, S., Pastushok, L., et al. (2024). hnRNP A1 dysfunction alters RNA splicing and drives neurodegeneration in multiple sclerosis (MS). Nature Communications 15, 356. 10.1038/s41467-023-44658-1.

28. Li, J., Guan, J., Qian, J., Feng, Y., Yao, R., Fan, R., and Wang, Z. (2018). Paean: A parallel transcriptome quantification tool combining gene expression and alternative splicing events using GPU. 3-6 Dec. 2018. pp. 676–679.

29. Zhu, Z., Zheng, Z., Zhang, F., Wu, Y., Trzaskowski, M., Maier, R., Robinson, M.R., McGrath, J.J., Visscher, P.M., Wray, N.R., and Yang, J. (2018). Causal associations between risk factors and common diseases inferred from GWAS summary data. Nature Communications 9, 224. 10.1038/s41467-017-02317-2.

30. Garrido-Martín, D., Borsari, B., Calvo, M., Reverter, F., and Guigó, R. (2021). Identification and analysis of splicing quantitative trait loci across multiple tissues in the human genome. Nature Communications 12, 727. 10.1038/s41467-020-20578-2.

31. Ezkurdia, I., Rodriguez, J.M., Carrillo-de Santa Pau, E., Vázquez, J., Valencia, A., and Tress, M.L. (2015). Most highly expressed protein-coding genes have a single dominant isoform. J Proteome Res 14, 1880–1887. 10.1021/pr501286b.

32. Single-cell transcriptomics of 20 mouse organs creates a Tabula Muris. (2018). Nature 562, 367–372. 10.1038/s41586-018-0590-4.

33. Harrison, D.E., Strong, R., Sharp, Z.D., Nelson, J.F., Astle, C.M., Flurkey, K., Nadon, N.L., Wilkinson, J.E., Frenkel, K., Carter, C.S., et al. (2009). Rapamycin fed late in life extends lifespan in genetically heterogeneous mice. Nature 460, 392–395. 10.1038/nature08221.

34. Mannick, J.B., Del Giudice, G., Lattanzi, M., Valiante, N.M., Praestgaard, J., Huang, B., Lonetto, M.A., Maecker, H.T., Kovarik, J., Carson, S., et al. (2014). mTOR inhibition improves immune function in the elderly. Sci Transl Med 6, 268ra179. 10.1126/scitranslmed.3009892.

35. Raien, A., Davis, S., Zhang, M., Zitser, D., Lin, M., Pitcher, G., Bhalodia, K., Subbian, S., and Venketaraman, V. (2023). Effects of Everolimus in Modulating the Host Immune Responses against Mycobacterium tuberculosis Infection. Cells 12, 2653.

36. Mannick, J.B., and Lamming, D.W. (2023). Targeting the biology of aging with mTOR inhibitors. Nature Aging 3, 642–660. 10.1038/s43587-023-00416-y.

37. Chang, J.W., Yeh, H.S., Park, M., Erber, L., Sun, J., Cheng, S., Bui, A.M., Fahmi, N.A., Nasti, R., Kuang, R., et al. (2019). mTOR-regulated U2af1 tandem exon splicing specifies transcriptome features for translational control. Nucleic Acids Res 47, 10373–10387. 10.1093/nar/gkz761.

38. Das, J.K., Banskota, N., Candia, J., Griswold, M.E., Orenduff, M., de Cabo, R., Corcoran, D.L., Das, S.K., De, S., Huffman, K.M., et al. (2023). Calorie restriction modulates the transcription of genes related to stress response and longevity in human muscle: The CALERIE study. Aging Cell 22, e13963. 10.1111/acel.13963.

39. López-Lluch, G., and Navas, P. (2016). Calorie restriction as an intervention in ageing. J Physiol 594, 2043–2060. 10.1113/jp270543.

40. Flanagan, E.W., Most, J., Mey, J.T., and Redman, L.M. (2020). Calorie Restriction and Aging in Humans. Annu Rev Nutr 40, 105–133. 10.1146/annurev-nutr-122319-034601.

41. Martin-Montalvo, A., Mercken, E.M., Mitchell, S.J., Palacios, H.H., Mote, P.L., Scheibye-Knudsen, M., Gomes, A.P., Ward, T.M., Minor, R.K., Blouin, M.J., et al. (2013). Metformin improves healthspan and lifespan in mice. Nat Commun 4, 2192. 10.1038/ncomms3192.

42. Cabreiro, F., Au, C., Leung, K.Y., Vergara-Irigaray, N., Cochemé, H.M., Noori, T., Weinkove, D., Schuster, E., Greene, N.D., and Gems, D. (2013). Metformin retards aging in C. elegans by altering microbial folate and methionine metabolism. Cell 153, 228–239. 10.1016/j.cell.2013.02.035.

43. Kucejova, B., Duarte, J., Satapati, S., Fu, X., Ilkayeva, O., Newgard, C.B., Brugarolas, J., and Burgess, S.C. (2016). Hepatic mTORC1 Opposes Impaired Insulin Action to Control Mitochondrial Metabolism in Obesity. Cell Rep 16, 508–519. 10.1016/j.celrep.2016.06.006.

44. Tyshkovskiy, A., Kholdina, D., Ying, K., Davitadze, M., Molière, A., Tongu, Y., Kasahara, T., Kats, L.M., Vladimirova, A., Moldakozhayev, A., et al. (2024). Transcriptomic Hallmarks of Mortality Reveal Universal and Specific Mechanisms of Aging, Chronic Disease, and Rejuvenation. bioRxiv, 2024.2007.2004.601982. 10.1101/2024.07.04.601982.

45. Khan, S.S., Singer, B.D., and Vaughan, D.E. (2017). Molecular and physiological manifestations and measurement of aging in humans. Aging Cell 16, 624–633. 10.1111/acel.12601.

46. Biamonti, G., Amato, A., Belloni, E., Di Matteo, A., Infantino, L., Pradella, D., and Ghigna, C. (2021). Alternative splicing in Alzheimer’s disease. Aging Clin Exp Res 33, 747–758. 10.1007/s40520-019-01360-x.

47. Lu, Y., Yue, D., Xie, J., Cheng, L., and Wang, X. (2022). Ontology Specific Alternative Splicing Changes in Alzheimer’s Disease. Front Genet 13, 926049. 10.3389/fgene.2022.926049.

48. Xiao, Y., Cai, G.P., Feng, X., Li, Y.J., Guo, W.H., Guo, Q., Huang, Y., Su, T., Li, C.J., Luo, X.H., et al. (2023). Splicing factor YBX1 regulates bone marrow stromal cell fate during aging. Embo j 42, e111762. 10.15252/embj.2022111762.

49. Deschênes, M., and Chabot, B. (2017). The emerging role of alternative splicing in senescence and aging. Aging Cell 16, 918–933. 10.1111/acel.12646.

50. Hong, J., Min, S., Yoon, G., and Lim, S.B. (2023). SRSF7 downregulation induces cellular senescence through generation of MDM2 variants. Aging (Albany NY) 15, 14591–14606. 10.18632/aging.205420.

51. Harrison, D.E., Strong, R., Sharp, Z.D., Nelson, J.F., Astle, C.M., Flurkey, K., Nadon, N.L., Wilkinson, J.E., Frenkel, K., Carter, C.S., et al. (2009). Rapamycin fed late in life extends lifespan in genetically heterogeneous mice. Nature 460, 392–395. 10.1038/nature08221.

52. The GTEx Consortium atlas of genetic regulatory effects across human tissues. (2020). Science 369, 1318–1330. 10.1126/science.aaz1776.

53. Horvath, S. (2013). DNA methylation age of human tissues and cell types. Genome Biology 14, 3156. 10.1186/gb-2013-14-10-r115.

54. Timmers, P.R., Mounier, N., Lall, K., Fischer, K., Ning, Z., Feng, X., Bretherick, A.D., Clark, D.W., Shen, X., Esko, T., et al. (2019). Genomics of 1 million parent lifespans implicates novel pathways and common diseases and distinguishes survival chances. Elife 8. 10.7554/eLife.39856.

55. Purcell, S., Neale, B., Todd-Brown, K., Thomas, L., Ferreira, M.A., Bender, D., Maller, J., Sklar, P., de Bakker, P.I., Daly, M.J., and Sham, P.C. (2007). PLINK: a tool set for whole-genome association and population-based linkage analyses. Am J Hum Genet 81, 559–575. 10.1086/519795.

56. Yang, J., Lee, S.H., Goddard, M.E., and Visscher, P.M. (2011). GCTA: a tool for genome-wide complex trait analysis. Am J Hum Genet 88, 76–82. 10.1016/j.ajhg.2010.11.011.

57. Yeo, G., and Burge, C.B. (2004). Maximum entropy modeling of short sequence motifs with applications to RNA splicing signals. J Comput Biol 11, 377–394. 10.1089/1066527041410418.

58. Yu, G., Wang, L.G., Han, Y., and He, Q.Y. (2012). clusterProfiler: an R package for comparing biological themes among gene clusters. Omics 16, 284–287. 10.1089/omi.2011.0118.

59. Hahn, O., Grönke, S., Stubbs, T.M., Ficz, G., Hendrich, O., Krueger, F., Andrews, S., Zhang, Q., Wakelam, M.J., Beyer, A., et al. (2017). Dietary restriction protects from age-associated DNA methylation and induces epigenetic reprogramming of lipid metabolism. Genome Biol 18, 56. 10.1186/s13059-017-1187-1.

60. Lee, H.J., Donati, A., Feliers, D., Sun, Y., Ding, Y., Madesh, M., Salmon, A.B., Ikeno, Y., Ross, C., O’Connor, C.L., et al. (2021). Chloride channel accessory 1 integrates chloride channel activity and mTORC1 in aging-related kidney injury. Aging Cell 20, e13407. 10.1111/acel.13407.

61. Shcherbakov, D., Juskeviciene, R., Cortés Sanchón, A., Brilkova, M., Rehrauer, H., Laczko, E., and Böttger, E.C. (2021). Mitochondrial Mistranslation in Brain Provokes a Metabolic Response Which Mitigates the Age-Associated Decline in Mitochondrial Gene Expression. Int J Mol Sci 22. 10.3390/ijms22052746.

62. Bundy, J.L., Vied, C., and Nowakowski, R.S. (2017). Sex differences in the molecular signature of the developing mouse hippocampus. BMC Genomics 18, 237. 10.1186/s12864-017-3608-7.

63. Han, G., Hong, S.H., Lee, S.J., Hong, S.P., and Cho, C. (2021). Transcriptome Analysis of Testicular Aging in Mice. Cells 10. 10.3390/cells10112895.

64. Börsch, A., Ham, D.J., Mittal, N., Tintignac, L.A., Migliavacca, E., Feige, J.N., Rüegg, M.A., and Zavolan, M. (2021). Molecular and phenotypic analysis of rodent models reveals conserved and species-specific modulators of human sarcopenia. Commun Biol 4, 194. 10.1038/s42003-021-01723-z.

65. Han, Y., Li, L.Z., Kastury, N.L., Thomas, C.T., Lam, M.P.Y., and Lau, E. (2021). Transcriptome features of striated muscle aging and predictability of protein level changes. Mol Omics 17, 796–808. 10.1039/d1mo00178g.

66. Shen, S., Park, J.W., Lu, Z.X., Lin, L., Henry, M.D., Wu, Y.N., Zhou, Q., and Xing, Y. (2014). rMATS: robust and flexible detection of differential alternative splicing from replicate RNA-Seq data. Proc Natl Acad Sci U S A 111, E5593–5601. 10.1073/pnas.1419161111.

67. Yilmaz, A., Peretz, M., Aharony, A., Sagi, I., and Benvenisty, N. (2018). Defining essential genes for human pluripotent stem cells by CRISPR-Cas9 screening in haploid cells. Nat Cell Biol 20, 610–619. 10.1038/s41556-018-0088-1.

68. Vasciaveo, A., Arriaga, J.M., de Almeida, F.N., Zou, M., Douglass, E.F., Picech, F., Shibata, M., Rodriguez-Calero, A., de Brot, S., Mitrofanova, A., et al. (2023). OncoLoop: A Network-Based Precision Cancer Medicine Framework. Cancer Discov 13, 386–409. 10.1158/2159-8290.Cd-22-0342.

69. Silic-Benussi, M., Sharova, E., Corradin, A., Urso, L., Raimondi, V., Cavallari, I., Buldini, B., Francescato, S., Minuzzo, S.A., D’Agostino, D.M., and Ciminale, V. (2023). Repurposing Verapamil to Enhance Killing of T-ALL Cells by the mTOR Inhibitor Everolimus. Antioxidants (Basel) 12. 10.3390/antiox12030625.

70. Martinez-Nunez, R.T., Wallace, A., Coyne, D., Jansson, L., Rush, M., Ennajdaoui, H., Katzman, S., Bailey, J., Deinhardt, K., Sanchez-Elsner, T., and Sanford, J.R. (2017). Modulation of nonsense mediated decay by rapamycin. Nucleic Acids Res 45, 3448–3459. 10.1093/nar/gkw1109.

71. Lu, J., Zhu, X., Shui, J.E., Xiong, L., Gierahn, T., Zhang, C., Wood, M., Hally, S., Love, J.C., Li, H., et al. (2021). Rho/SMAD/mTOR triple inhibition enables long-term expansion of human neonatal tracheal aspirate-derived airway basal cell-like cells. Pediatr Res 89, 502–509. 10.1038/s41390-020-0925-3.

72. Nguyen, T.U., Hector, H., Pederson, E.N., Lin, J., Ouyang, Z., Wendel, H.G., and Singh, K. (2023). Rapamycin-Induced Feedback Activation of eIF4E-EIF4A Dependent mRNA Translation in Pancreatic Cancer. Cancers (Basel) 15. 10.3390/cancers15051444.

73. Golden, E., Rashwan, R., Woodward, E.A., Sgro, A., Wang, E., Sorolla, A., Waryah, C., Tie, W.J., Cuyàs, E., Ratajska, M., et al. (2021). The oncogene AAMDC links PI3K-AKT-mTOR signaling with metabolic reprograming in estrogen receptor-positive breast cancer. Nat Commun 12, 1920. 10.1038/s41467-021-22101-7.

74. Meng, Y., Xiang, R., Yan, H., Zhou, Y., Hu, Y., Yang, J., Zhou, Y., and Cui, Q. (2020). Transcriptomic landscape profiling of metformin-treated healthy mice: Implication for potential hypertension risk when prophylactically used. J Cell Mol Med 24, 8138–8150. 10.1111/jcmm.15472.

75. Li, B., and Dewey, C.N. (2011). RSEM: accurate transcript quantification from RNA-Seq data with or without a reference genome. BMC Bioinformatics 12, 323. 10.1186/1471-2105-12-323.

76. Seiler, M., Peng, S., Agrawal, A.A., Palacino, J., Teng, T., Zhu, P., Smith, P.G., Buonamici, S., and Yu, L. (2018). Somatic Mutational Landscape of Splicing Factor Genes and Their Functional Consequences across 33 Cancer Types. Cell Rep 23, 282–296.e284. 10.1016/j.celrep.2018.01.088.

77. Yeo, G.W., Coufal, N.G., Liang, T.Y., Peng, G.E., Fu, X.D., and Gage, F.H. (2009). An RNA code for the FOX2 splicing regulator revealed by mapping RNA-protein interactions in stem cells. Nat Struct Mol Biol 16, 130–137. 10.1038/nsmb.1545.

78. Lovci, M.T., Ghanem, D., Marr, H., Arnold, J., Gee, S., Parra, M., Liang, T.Y., Stark, T.J., Gehman, L.T., Hoon, S., et al. (2013). Rbfox proteins regulate alternative mRNA splicing through evolutionarily conserved RNA bridges. Nat Struct Mol Biol 20, 1434–1442. 10.1038/nsmb.2699.

79. Tyshkovskiy, A., Bozaykut, P., Borodinova, A.A., Gerashchenko, M.V., Ables, G.P., Garratt, M., Khaitovich, P., Clish, C.B., Miller, R.A., and Gladyshev, V.N. (2019). Identification and Application of Gene Expression Signatures Associated with Lifespan Extension. Cell Metab 30, 573–593.e578. 10.1016/j.cmet.2019.06.018.

80. Subramanian, A., Tamayo, P., Mootha, V.K., Mukherjee, S., Ebert, B.L., Gillette, M.A., Paulovich, A., Pomeroy, S.L., Golub, T.R., Lander, E.S., and Mesirov, J.P. (2005). Gene set enrichment analysis: a knowledge-based approach for interpreting genome-wide expression profiles. Proc Natl Acad Sci U S A 102, 15545–15550. 10.1073/pnas.0506580102.

