## Supplementary material for "Mammalian aging involves genome-wide splicing degeneration leading to functional decline": SuppleFigures

### Figure S1

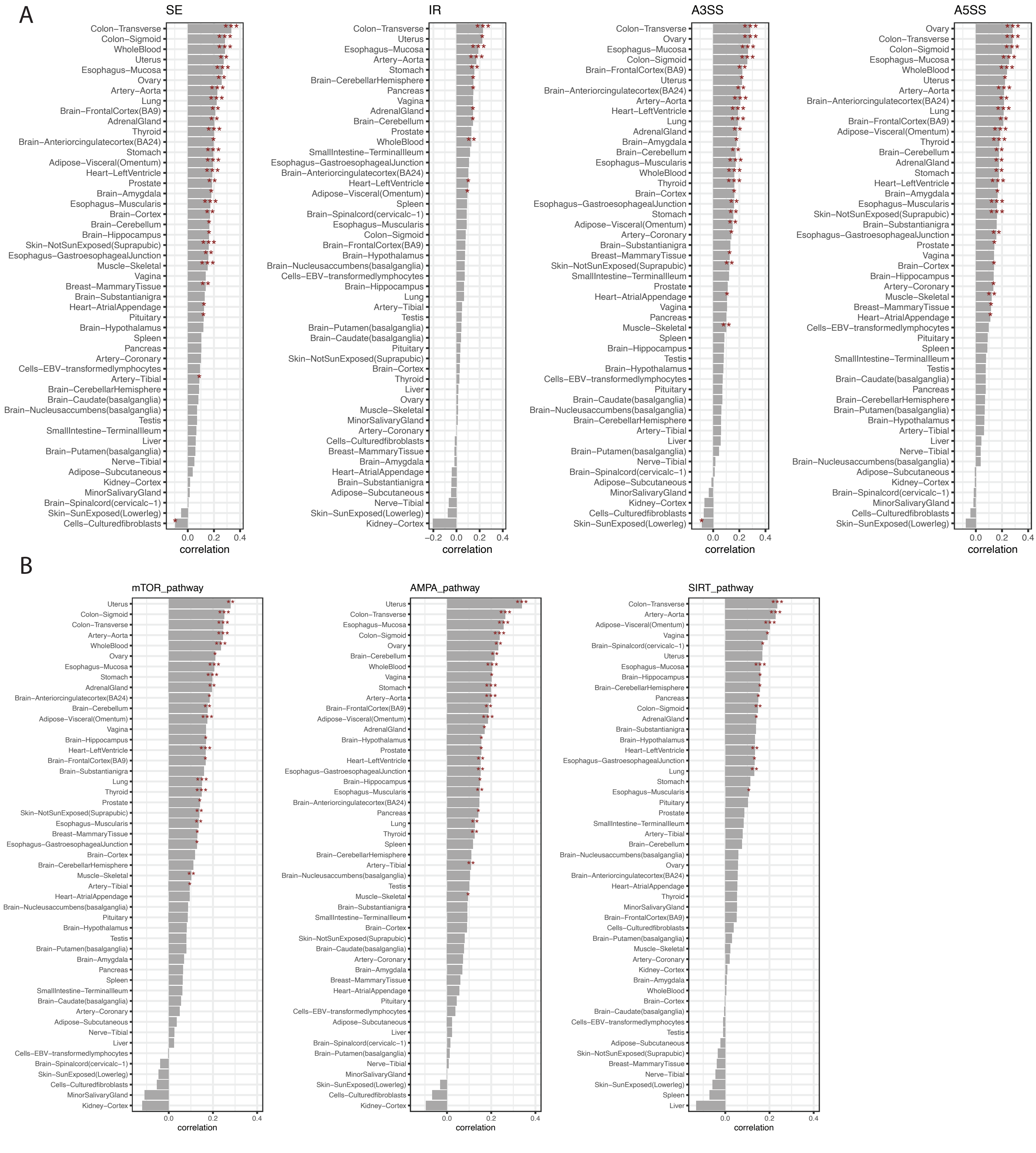

A
SE
IR
A3SS
A5SS
correlation
B
mTOR_pathway
AMPA_pathway
SIRT_pathway
correlation

### Figure S2

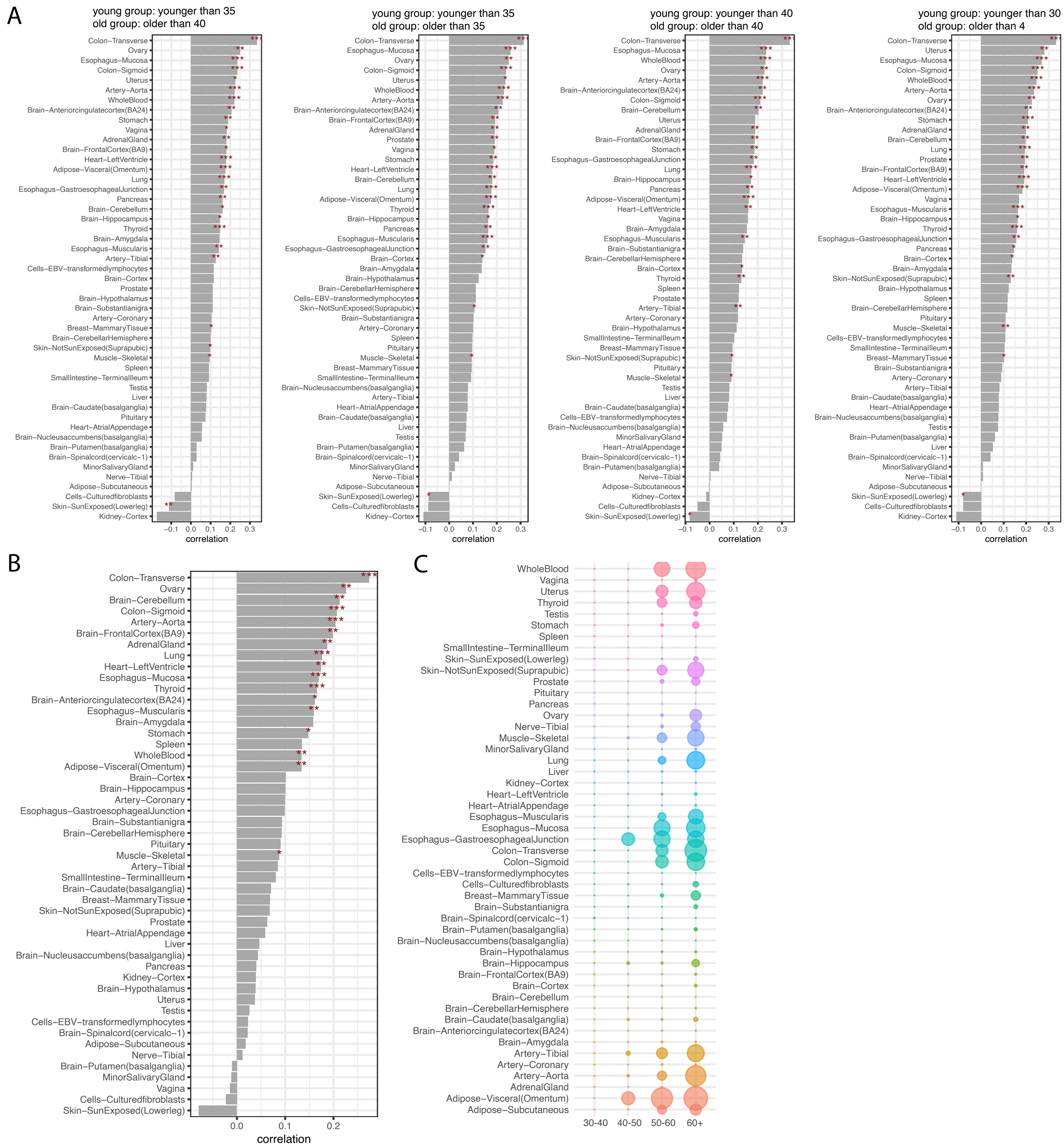
A
young group: younger than 35
old group: older than 40
young group: younger than 35
old group: older than 35
young group: younger than 40
old group: older than 40
young group: younger than 30
old group: older than 4
correlation
B
correlation
C
30-40
40-50
50-60
60+

### Figure S3

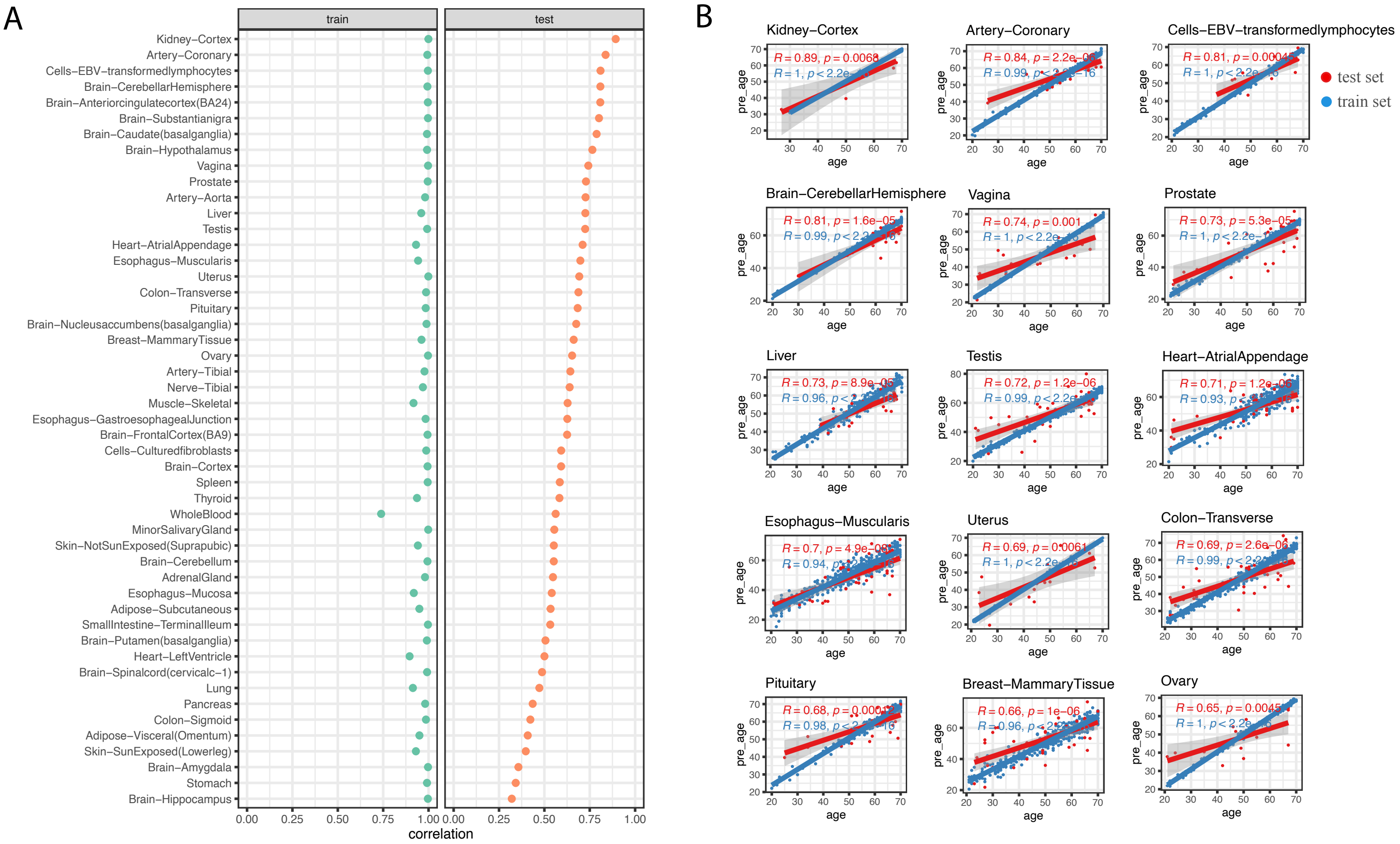
A
train
test
Kidney−Cortex
Artery−Coronary
Cells−EBV−transformedlymphocytes
Brain−CerebellarHemisphere
Brain−Anteriorcingulatecortex(BA24)
Brain−Substantianigra
Brain−Caudate(basalganglia)
Brain−Hypothalamus
Vagina
Prostate
Artery−Aorta
Liver
Testis
Heart−AtrialAppendage
Esophagus−Muscularis
Uterus
Colon−Transverse
Pituitary
Brain−Nucleusaccumbens(basalganglia)
Breast−MammaryTissue
Ovary
Artery−Tibial
Nerve−Tibial
Muscle−Skeletal
Esophagus−GastroesophagealJunction
Brain−FrontalCortex(BA9)
Cells−Culturedfibroblasts
Brain−Cortex
Spleen
Thyroid
WholeBlood
MinorSalivaryGland
Skin−NotSunExposed(Suprapubic)
Brain−Cerebellum
AdrenalGland
Esophagus−Mucosa
Adipose−Subcutaneous
SmallIntestine−TerminalIleum
Brain−Putamen(basalganglia)
Heart−LeftVentricle
Brain−Spinalcord(cervicalc−1)
Lung
Pancreas
Colon−Sigmoid
Adipose−Visceral(Omentum)
Skin−SunExposed(Lowerleg)
Brain−Amygdala
Stomach
Brain−Hippocampus
0.00 0.25 0.50 0.75 1.00
correlation
B
test set
train set
Kidney−Cortex
R = 0.89, p = 0.0068
Artery−Coronary
R = 0.84, p = 2.2e−06
Cells−EBV−transformedlymphocytes
R = 0.81, p = 0.00045
Brain−CerebellarHemisphere
R = 0.81, p = 1.6e−05
Vagina
R = 0.74, p = 0.001
Prostate
R = 0.73, p = 5.3e−05
Liver
Testis
R = 0.72, p = 1.2e−06
Heart−AtrialAppendage
Esophagus−Muscularis
Uterus
R = 0.69, p = 0.0061
Colon−Transverse
R = 0.69, p = 2.6e−06
Pituitary
Breast−MammaryTissue
R = 0.66, p = 1e−06
Ovary
pre_age
age

### Figure S4

A

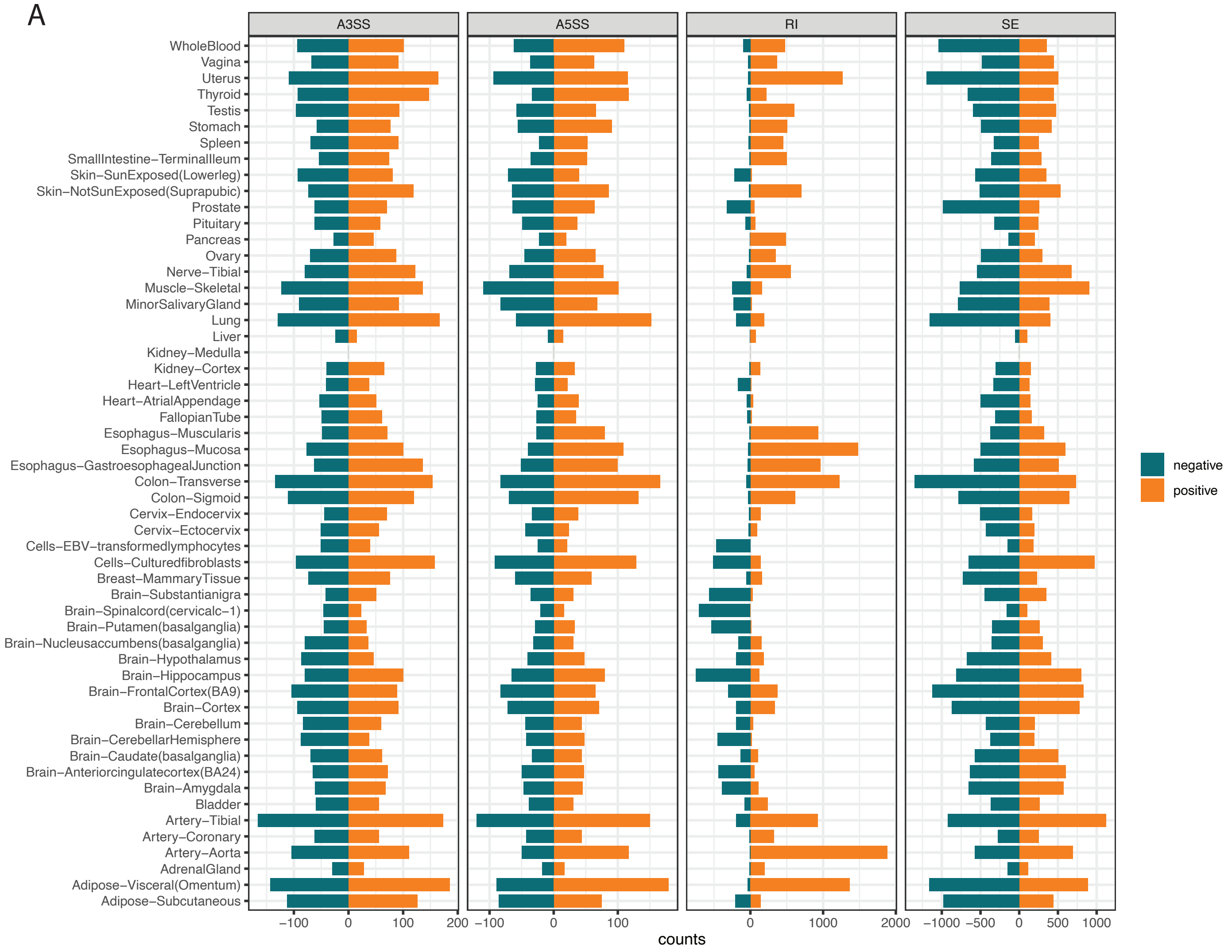

A3SS
A5SS
RI
SE
WholeBlood
Vagina
Uterus
Thyroid
Testis
Stomach
Spleen
SmallIntestine–TerminalIleum
Skin–SunExposed(Lowerleg)
Skin–NotSunExposed(Suprapubic)
Prostate
Pituitary
Pancreas
Ovary
Nerve–Tibial
Muscle–Skeletal
MinorSalivaryGland
Lung
Liver
Kidney–Medulla
Kidney–Cortex
Heart–LeftVentricle
Heart–AtrialAppendage
FallopianTube
Esophagus–Muscularis
Esophagus–Mucosa
Esophagus–GastroesophagealJunction
Colon–Transverse
Colon–Sigmoid
Cervix–Endocervix
Cervix–Ectocervix
Cells–EBV–transformedlymphocytes
Cells–Culturedfibroblasts
Breast–MammaryTissue
Brain–Substantianigra
Brain–Spinalcord(cervicalc–1)
Brain–Putamen(basalganglia)
Brain–Nucleusaccumbens(basalganglia)
Brain–Hypothalamus
Brain–Hippocampus
Brain–FrontalCortex(BA9)
Brain–Cortex
Brain–Cerebellum
Brain–CerebellarHemisphere
Brain–Caudate(basalganglia)
Brain–Anteriorcingulatecortex(BA24)
Brain–Amygdala
Bladder
Artery–Tibial
Artery–Coronary
Artery–Aorta
AdrenalGland
Adipose–Visceral(Omentum)
Adipose–Subcutaneous
–100
0
100
200
–100
0
100
0
1000
2000
–1000
–500
0
500
1000
counts
negative
positive

### Figure S5

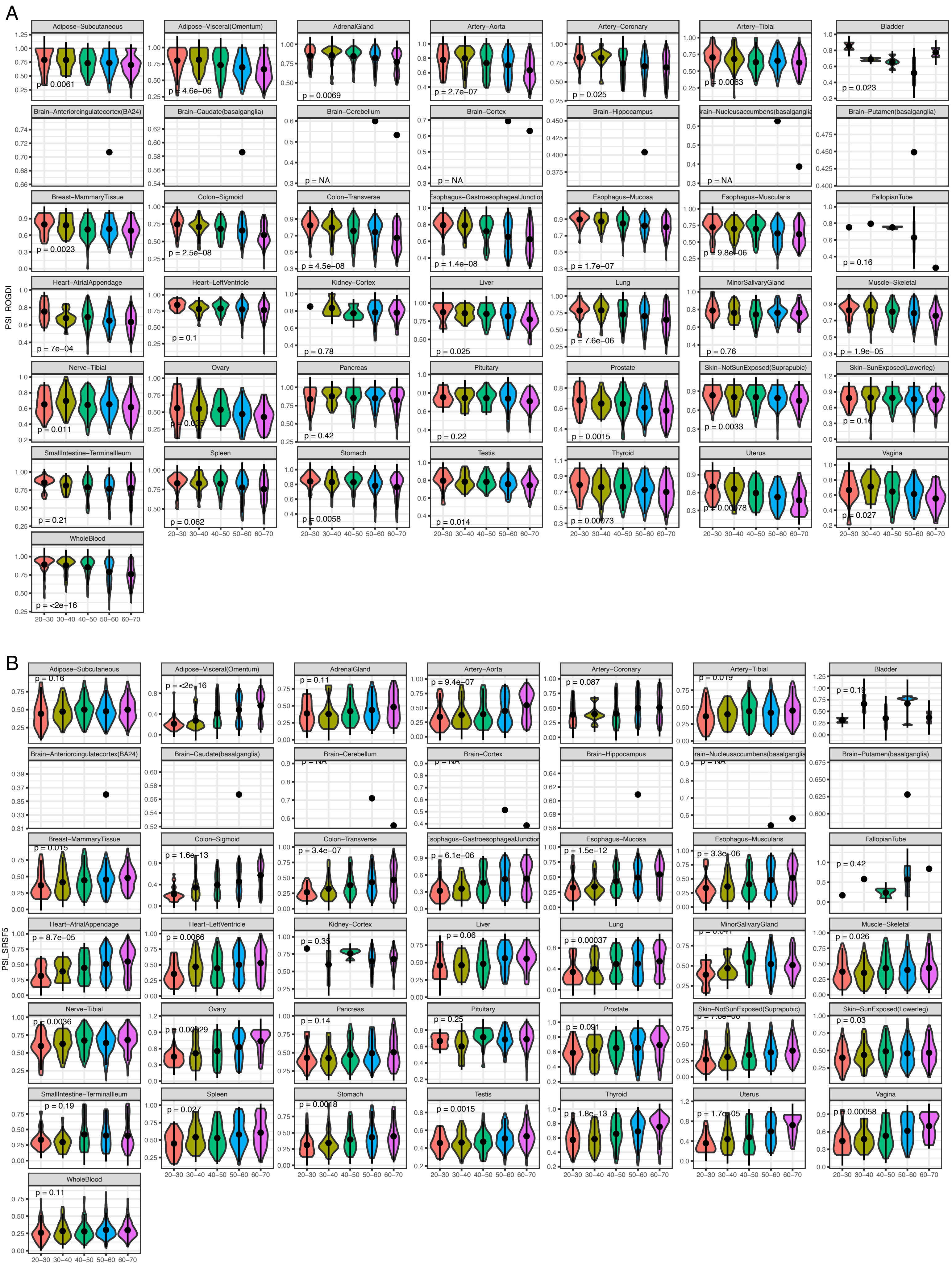
A
Adipose−Subcutaneous
p = 0.0061
Adipose−Visceral(Omentum)
p = 4.6e−06
AdrenalGland
p = 0.0069
Artery−Aorta
p = 2.7e−07
Artery−Coronary
p = 0.025
Artery−Tibial
p = 0.0063
Bladder
p = 0.023
Brain−Anteriorcingulatecortex(BA24)
Brain−Caudate(basalganglia)
Brain−Cerebellum
p = NA
Brain−Cortex
p = NA
Brain−Hippocampus
Brain−Putamen(basalganglia)
p = NA
Breast−MammaryTissue
p = 0.0023
Colon−Sigmoid
p = 2.5e−08
Colon−Transverse
p = 4.5e−08
p = 1.4e−08
Esophagus−Mucosa
p = 1.7e−07
Esophagus−Muscularis
p = 9.8e−06
FallopianTube
p = 0.16
Heart−AtrialAppendage
p = 7e−04
Heart−LeftVentricle
p = 0.1
Kidney−Cortex
p = 0.78
Liver
p = 0.025
Lung
p = 7.6e−06
MinorSalivaryGland
p = 0.76
Muscle−Skeletal
p = 1.9e−05
Nerve−Tibial
p = 0.011
Ovary
Pancreas
p = 0.42
Pituitary
p = 0.22
Prostate
p = 0.0015
Skin−NotSunExposed(Suprapubic)
p = 0.0033
Skin−SunExposed(Lowerleg)
SmallIntestine−TerminalIleum
p = 0.21
Spleen
p = 0.062
Stomach
p = 0.0058
Testis
p = 0.014
Thyroid
p = 0.00073
Uterus
Vagina
p = 0.027
WholeBlood
p = <2e−16
PSI_ROGDI
20−30 30−40 40−50 50−60 60−70
B
Adipose−Subcutaneous
p = 0.16
Adipose−Visceral(Omentum)
p = <2e−16
AdrenalGland
p = 0.11
Artery−Aorta
p = 9.4e−07
Artery−Coronary
p = 0.087
Artery−Tibial
p = 0.019
Bladder
p = 0.19
Brain−Anteriorcingulatecortex(BA24)
Brain−Caudate(basalganglia)
Brain−Cerebellum
Brain−Cortex
Brain−Hippocampus
Brain−Putamen(basalganglia)
Breast−MammaryTissue
p = 0.015
Colon−Sigmoid
p = 1.6e−13
Colon−Transverse
p = 3.4e−07
p = 6.1e−06
Esophagus−Mucosa
p = 1.5e−12
Esophagus−Muscularis
p = 3.3e−06
FallopianTube
p = 0.42
Heart−AtrialAppendage
p = 8.7e−05
Heart−LeftVentricle
p = 0.0066
Kidney−Cortex
p = 0.35
Liver
p = 0.06
Lung
p = 0.00037
MinorSalivaryGland
Muscle−Skeletal
p = 0.026
Nerve−Tibial
p = 0.0036
Ovary
Pancreas
p = 0.14
Pituitary
p = 0.25
Prostate
p = 0.091
Skin−NotSunExposed(Suprapubic)
Skin−SunExposed(Lowerleg)
p = 0.03
SmallIntestine−TerminalIleum
p = 0.19
Spleen
p = 0.027
Stomach
p = 0.0018
Testis
p = 0.0015
Thyroid
p = 1.8e−13
Uterus
p = 1.7e−05
Vagina
p = 0.00058
WholeBlood
p = 0.11
PSI_SRSF5
20−30 30−40 40−50 50−60 60−70

### Figure S6

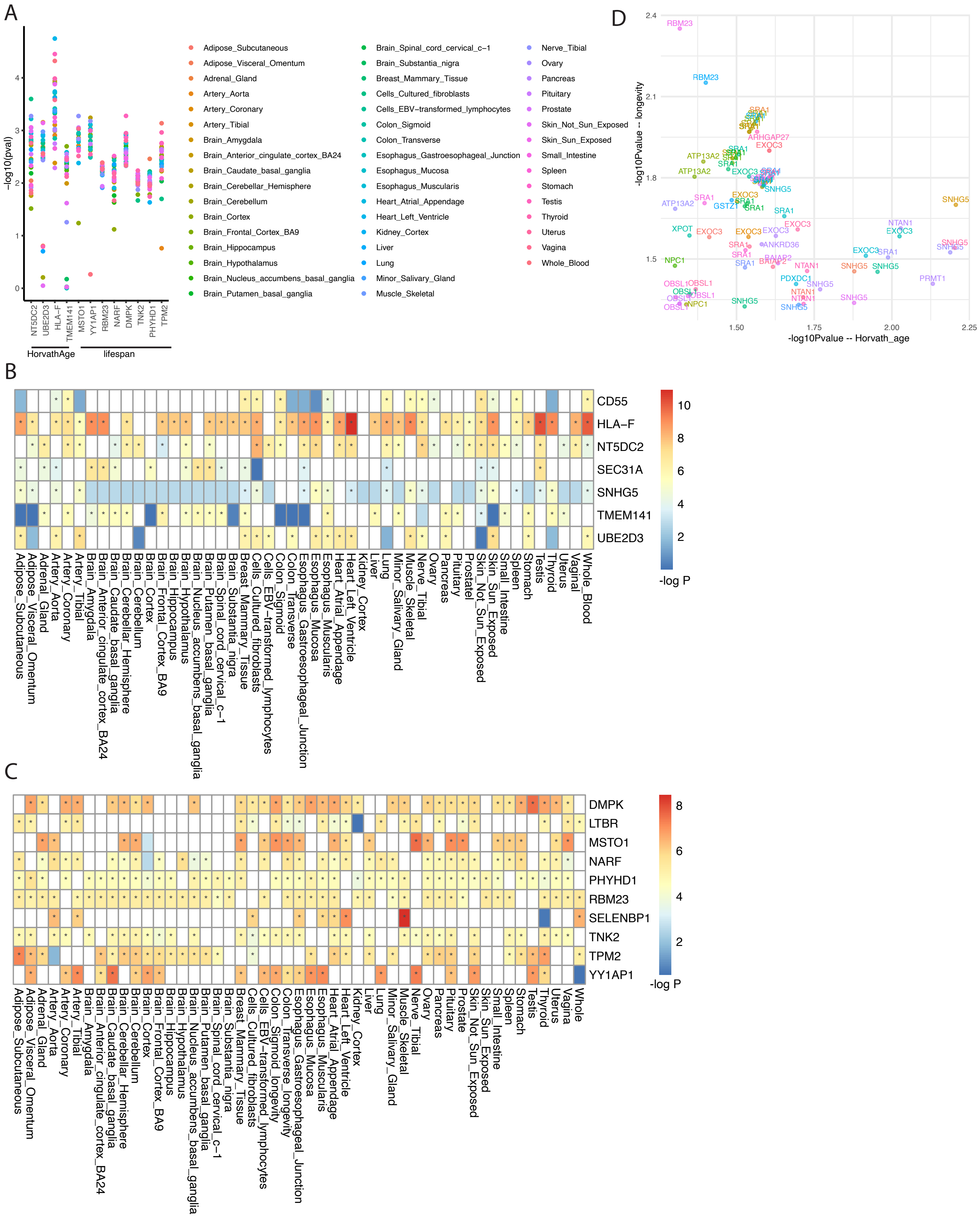
A
-log10(pval)
NT5DC2
UBE2D3
HLA−F
TMEM141
MSTO1
YY1AP1
RBM23
NARF
DMPK
TNK2
PHYHD1
TPM2
HorvathAge
lifespan
Adipose_Subcutaneous
Adipose_Visceral_Omentum
Adrenal_Gland
Artery_Aorta
Artery_Coronary
Artery_Tibial
Brain_Amygdala
Brain_Anterior_cingulate_cortex_BA24
Brain_Caudate_basal_ganglia
Brain_Cerebellar_Hemisphere
Brain_Cerebellum
Brain_Cortex
Brain_Frontal_Cortex_BA9
Brain_Hippocampus
Brain_Hypothalamus
Brain_Nucleus_accumbens_basal_ganglia
Brain_Putamen_basal_ganglia
Brain_Spinal_cord_cervical_c−1
Brain_Substantia_nigra
Breast_Mammary_Tissue
Cells_Cultured_fibroblasts
Cells_EBV−transformed_lymphocytes
Colon_Sigmoid
Colon_Transverse
Esophagus_Gastroesophageal_Junction
Esophagus_Mucosa
Esophagus_Muscularis
Heart_Atrial_Appendage
Heart_Left_Ventricle
Kidney_Cortex
Liver
Lung
Minor_Salivary_Gland
Muscle_Skeletal
Nerve_Tibial
Ovary
Pancreas
Pituitary
Prostate
Skin_Not_Sun_Exposed
Skin_Sun_Exposed
Small_Intestine
Spleen
Stomach
Testis
Thyroid
Uterus
Vagina
Whole_Blood
D
-log10Pvalue -- longevity
-log10Pvalue -- Horvath_age
B
CD55
HLA−F
NT5DC2
SEC31A
SNHG5
TMEM141
UBE2D3
-log P
C
DMPK
LTBR
MSTO1
NARF
PHYHD1
RBM23
SELENBP1
TNK2
TPM2
YY1AP1
-log P

### Figure S7

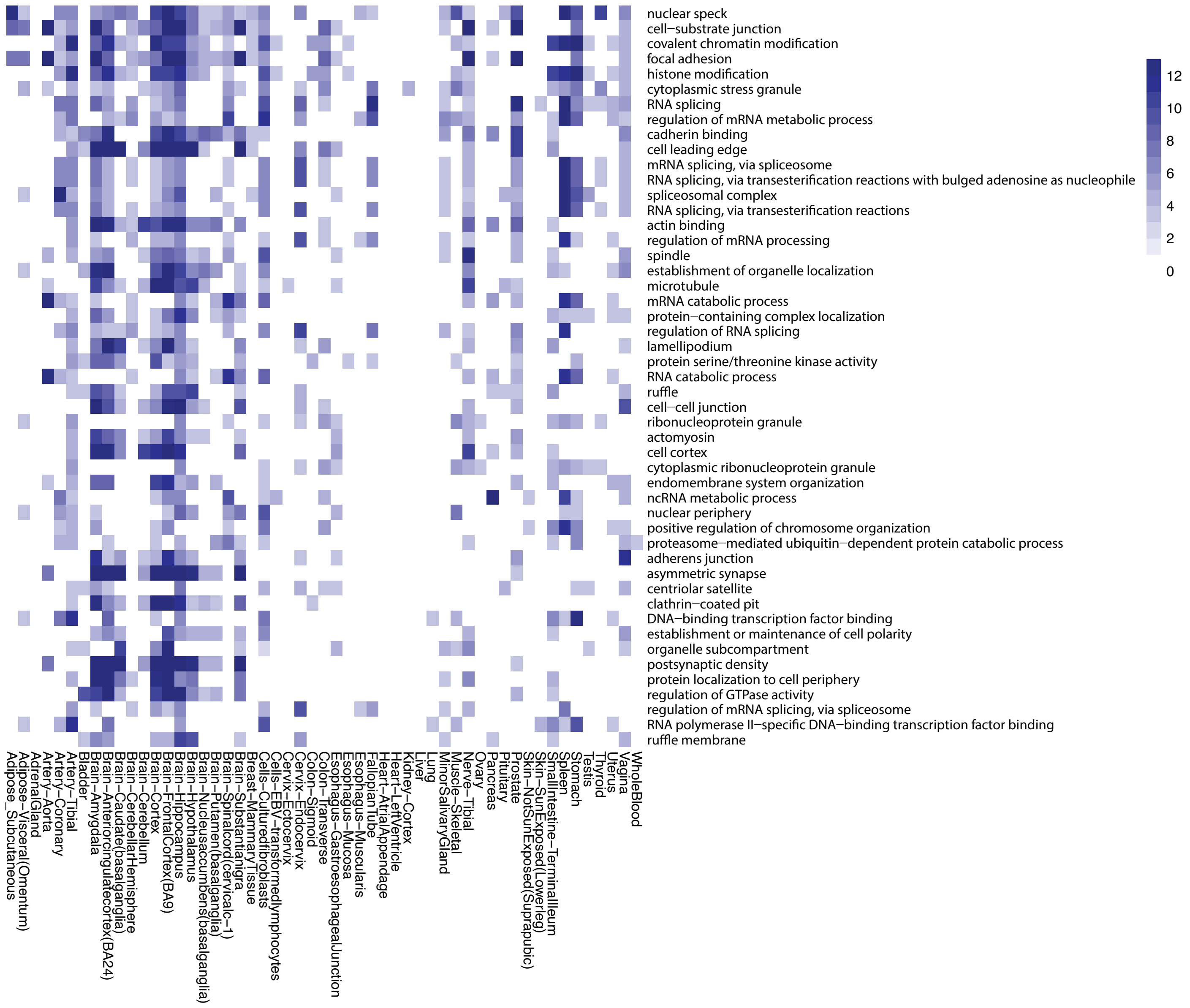
nuclear speck
cell−substrate junction
covalent chromatin modification
focal adhesion
histone modification
cytoplasmic stress granule
RNA splicing
regulation of mRNA metabolic process
cadherin binding
cell leading edge
mRNA splicing, via spliceosome
RNA splicing, via transesterification reactions with bulged adenosine as nucleophile
spliceosomal complex
RNA splicing, via transesterification reactions
actin binding
regulation of mRNA processing
spindle
establishment of organelle localization
microtubule
mRNA catabolic process
protein−containing complex localization
regulation of RNA splicing
lamellipodium
protein serine/threonine kinase activity
RNA catabolic process
ruffle
cell−cell junction
ribonucleoprotein granule
actomyosin
cell cortex
cytoplasmic ribonucleoprotein granule
endomembrane system organization
ncRNA metabolic process
nuclear periphery
positive regulation of chromosome organization
proteasome−mediated ubiquitin−dependent protein catabolic process
adherens junction
asymmetric synapse
centriolar satellite
clathrin−coated pit
DNA−binding transcription factor binding
establishment or maintenance of cell polarity
organelle subcompartment
postsynaptic density
protein localization to cell periphery
regulation of GTPase activity
regulation of mRNA splicing, via spliceosome
RNA polymerase II−specific DNA−binding transcription factor binding
ruffle membrane
Adipose_Subcutaneous
Adipose−Visceral(Omentum)
AdrenalGland
Artery−Aorta
Artery−Coronary
Artery−Tibial
Bladder
Brain−Amygdala
Brain−Anteriorcingulatecortex(BA24)
Brain−Caudate(basalganglia)
Brain−CerebellarHemisphere
Brain−Cerebellum
Brain−Cortex
Brain−FrontalCortex(BA9)
Brain−Hippocampus
Brain−Hypothalamus
Brain−Nucleusaccumbens(basalganglia)
Brain−Putamen(basalganglia)
Brain−Spinalcord(cervicalc-1)
Brain−Substantianigra
Breast−MammaryTissue
Cells−Culturedfibroblasts
Cells−EBV−transformedlymphocytes
Cervix−Ectocervix
Cervix−Endocervix
Colon−Sigmoid
Colon−Transverse
Esophagus−GastroesophagealJunction
Esophagus−Mucosa
Esophagus−Muscularis
FallopianTube
Heart−AtrialAppendage
Heart−LeftVentricle
Kidney−Cortex
Liver
Lung
MinorSalivaryGland
Muscle−Skeletal
Nerve−Tibial
Ovary
Pancreas
Pituitary
Prostate
Skin−NotSunExposed(Suprapubic)
Skin−SunExposed(Lowerleg)
SmallIntestine−TerminalIleum
Spleen
Stomach
Testis
Thyroid
Uterus
Vagina
WholeBlood
12
10
8
6
4
2
0

### Figure S8

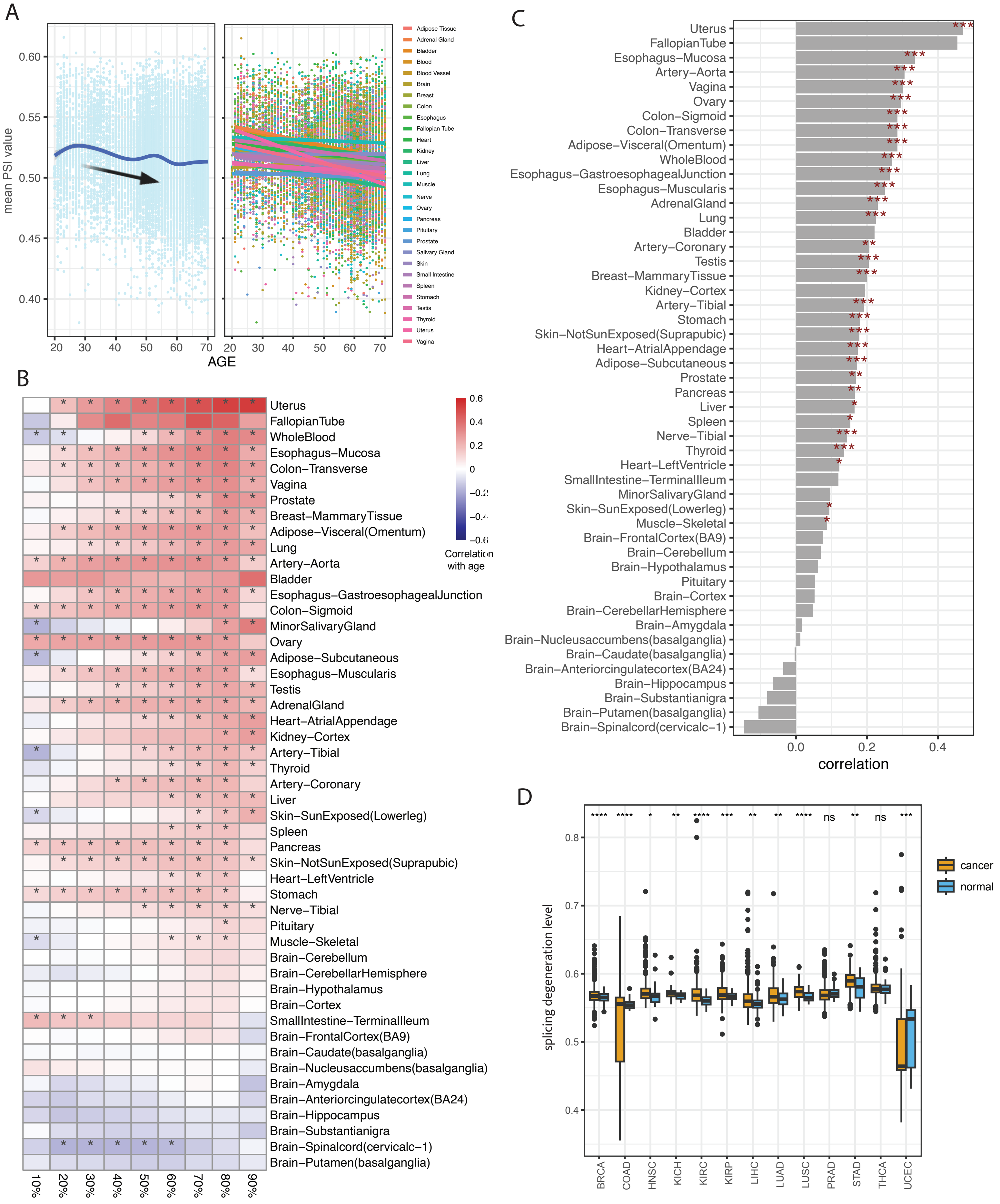
A
mean PSI value
0.60
0.55
0.50
0.45
0.40
20 30 40 50 60 70
AGE
Adipose Tissue
Adrenal Gland
Bladder
Blood
Blood Vessel
Brain
Breast
Colon
Esophagus
Fallopian Tube
Heart
Kidney
Liver
Lung
Muscle
Nerve
Ovary
Pancreas
Pituitary
Prostate
Salivary Gland
Skin
Small Intestine
Spleen
Stomach
Testis
Thyroid
Uterus
Vagina
B
Uterus
FallopianTube
WholeBlood
Esophagus–Mucosa
Colon–Transverse
Vagina
Prostate
Breast–MammaryTissue
Adipose–Visceral(Omentum)
Lung
Artery–Aorta
Bladder
Esophagus–GastroesophagealJunction
Colon–Sigmoid
MinorSalivaryGland
Ovary
Adipose–Subcutaneous
Esophagus–Muscularis
Testis
AdrenalGland
Heart–AtrialAppendage
Kidney–Cortex
Artery–Tibial
Thyroid
Artery–Coronary
Liver
Skin–SunExposed(Lowerleg)
Spleen
Pancreas
Skin–NotSunExposed(Suprapubic)
Heart–LeftVentricle
Stomach
Nerve–Tibial
Pituitary
Muscle–Skeletal
Brain–Cerebellum
Brain–CerebellarHemisphere
Brain–Hypothalamus
Brain–Cortex
SmallIntestine–TerminalIleum
Brain–FrontalCortex(BA9)
Brain–Caudate(basalganglia)
Brain–Nucleusaccumbens(basalganglia)
Brain–Amygdala
Brain–Anteriorcingulatecortex(BA24)
Brain–Hippocampus
Brain–Substantianigra
Brain–Spinalcord(cervicalc–1)
Brain–Putamen(basalganglia)
10% 20% 30% 40% 50% 60% 70% 80% 90%
0.6
0.4
0.2
0
Correlation with age
C
Uterus
FallopianTube
Esophagus–Mucosa
Artery–Aorta
Vagina
Ovary
Colon–Sigmoid
Colon–Transverse
Adipose–Visceral(Omentum)
WholeBlood
Esophagus–GastroesophagealJunction
Esophagus–Muscularis
AdrenalGland
Lung
Bladder
Artery–Coronary
Testis
Breast–MammaryTissue
Kidney–Cortex
Artery–Tibial
Stomach
Skin–NotSunExposed(Suprapubic)
Heart–AtrialAppendage
Adipose–Subcutaneous
Prostate
Pancreas
Liver
Spleen
Nerve–Tibial
Thyroid
Heart–LeftVentricle
SmallIntestine–TerminalIleum
MinorSalivaryGland
Skin–SunExposed(Lowerleg)
Muscle–Skeletal
Brain–FrontalCortex(BA9)
Brain–Cerebellum
Brain–Hypothalamus
Pituitary
Brain–Cortex
Brain–CerebellarHemisphere
Brain–Amygdala
Brain–Nucleusaccumbens(basalganglia)
Brain–Caudate(basalganglia)
Brain–Anteriorcingulatecortex(BA24)
Brain–Hippocampus
Brain–Substantianigra
Brain–Putamen(basalganglia)
Brain–Spinalcord(cervicalc–1)
0.0 0.2 0.4
correlation
D
splicing degeneration level
0.8
0.7
0.6
0.5
0.4
**** **** * ** **** *** ** ** **** ns ** ns ***
BRCA COAD HNSC KICH KIRC KIRP LIHC LUAD LUSC PRAD STAD THCA UCEC
cancer
normal

### Figure S9

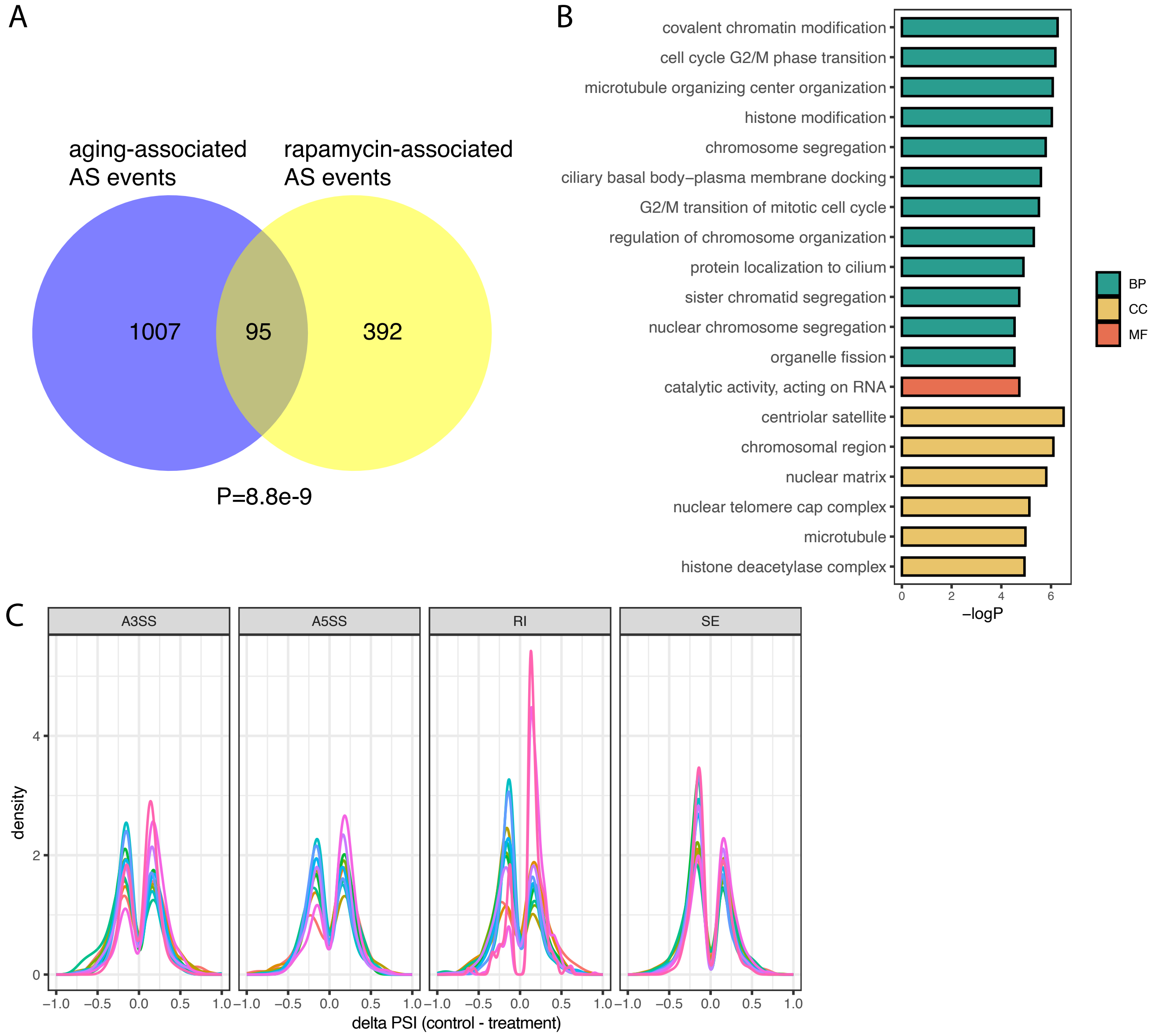
A
aging-associated AS events
rapamycin-associated AS events
1007
95
392
P=8.8e-9
B
covalent chromatin modification
cell cycle G2/M phase transition
microtubule organizing center organization
histone modification
chromosome segregation
ciliary basal body–plasma membrane docking
G2/M transition of mitotic cell cycle
regulation of chromosome organization
protein localization to cilium
sister chromatid segregation
nuclear chromosome segregation
organelle fission
catalytic activity, acting on RNA
centriolar satellite
chromosomal region
nuclear matrix
nuclear telomere cap complex
microtubule
histone deacetylase complex
0
2
4
6
−logP
BP
CC
MF
C
A3SS
A5SS
RI
SE
density
0
2
4
−1.0
−0.5
0.0
0.5
1.0
delta PSI (control - treatment)

### Figure S10

A

$$YTPM = \beta^0 + \beta^1 AGE + \beta^2\ GENDER + \beta^3\ RACE + \beta^4\ DTHHRDY + \varepsilon$$

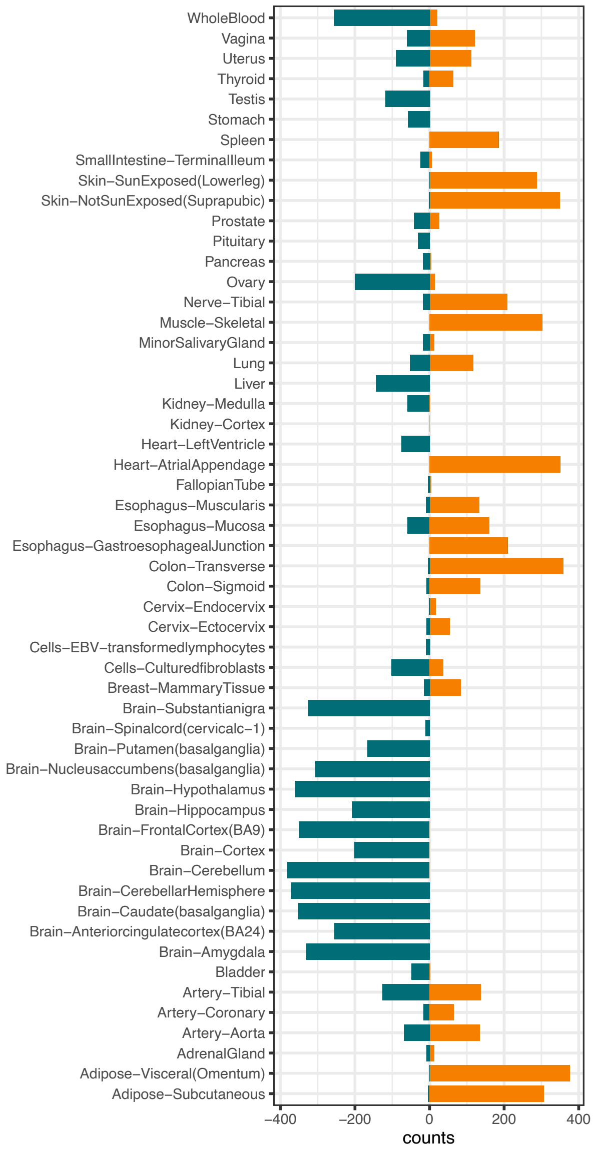


B

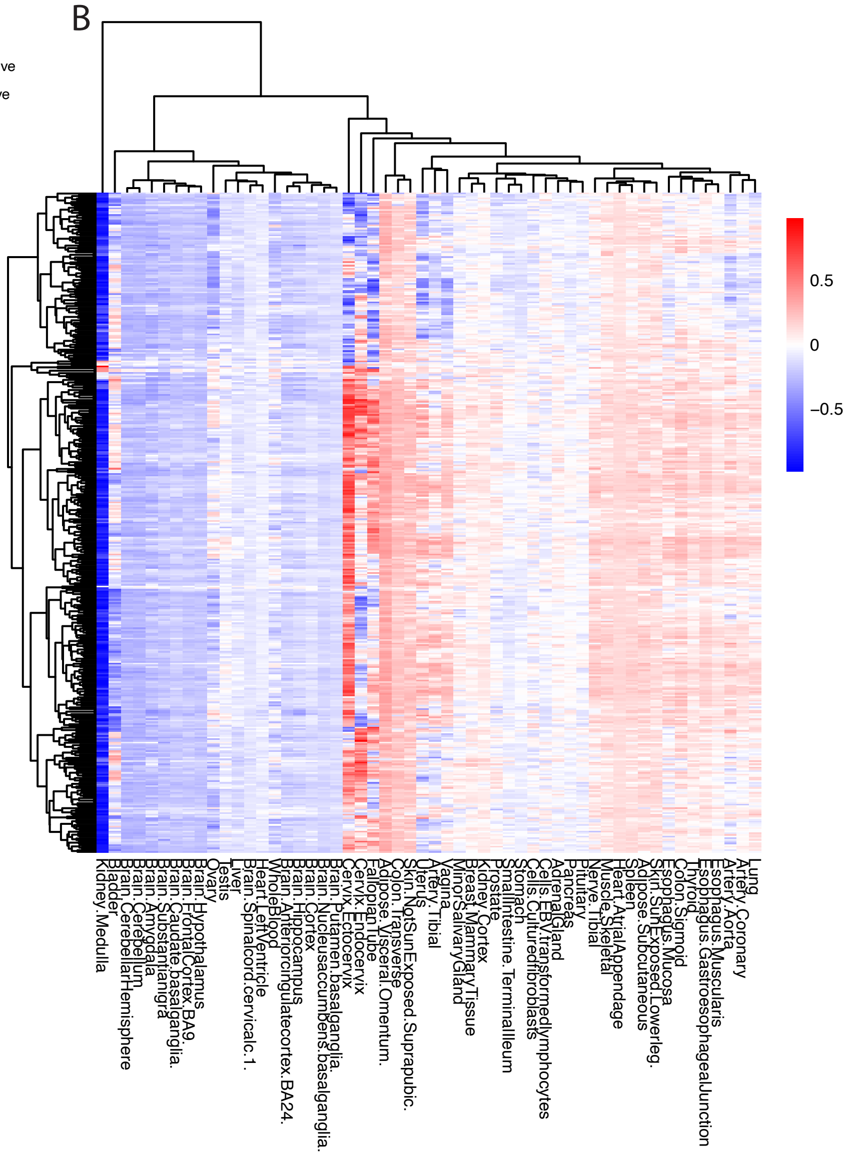

Figure S11

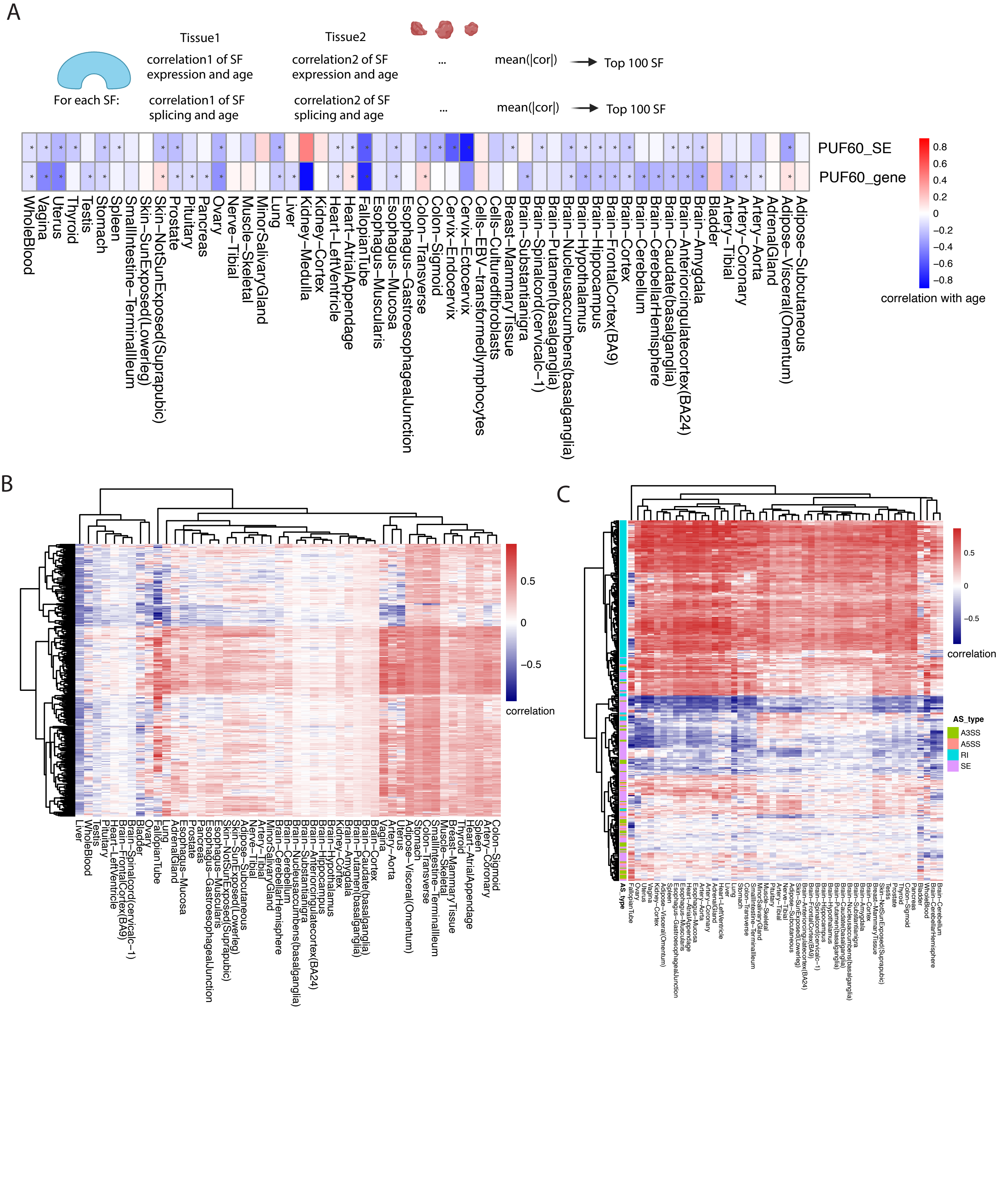
A
Tissue1
Tissue2
For each SF:
correlation1 of SF expression and age
correlation2 of SF expression and age
...
mean(|cor|)
Top 100 SF
correlation1 of SF splicing and age
correlation2 of SF splicing and age
...
mean(|cor|)
Top 100 SF
PUF60_SE
PUF60_gene
correlation with age
WholeBlood
Vagina
Uterus
Thyroid
Testis
Stomach
Spleen
SmallIntestine−TerminalIleum
Skin−SunExposed(Lowerleg)
Skin−NotSunExposed(Suprapubic)
Prostate
Pituitary
Pancreas
Ovary
Nerve−Tibial
Muscle−Skeletal
MinorSalivaryGland
Lung
Liver
Kidney−Medulla
Kidney−Cortex
Heart−LeftVentricle
Heart−AtrialAppendage
FallopianTube
Esophagus−Muscularis
Esophagus−Mucosa
Esophagus−GastroesophagealJunction
Colon−Transverse
Colon−Sigmoid
Cervix−Endocervix
Cervix−Ectocervix
Cells−EBV−transformedlymphocytes
Cells−Culturedfibroblasts
Breast−MammaryTissue
Brain−Substantianigra
Brain−Spinalcord(cervicalc−1)
Brain−Putamen(basalganglia)
Brain−Nucleusaccumbens(basalganglia)
Brain−Hypothalamus
Brain−Hippocampus
Brain−FrontalCortex(BA9)
Brain−Cortex
Brain−Cerebellum
Brain−CerebellarHemisphere
Brain−Caudate(basalganglia)
Brain−Anteriorcingulatecortex(BA24)
Brain−Amygdala
Bladder
Artery−Tibial
Artery−Coronary
Artery−Aorta
AdrenalGland
Adipose−Visceral(Omentum)
Adipose−Subcutaneous
B
correlation
C
correlation
AS_type
A3SS
A5SS
RI
SE

Figure S12

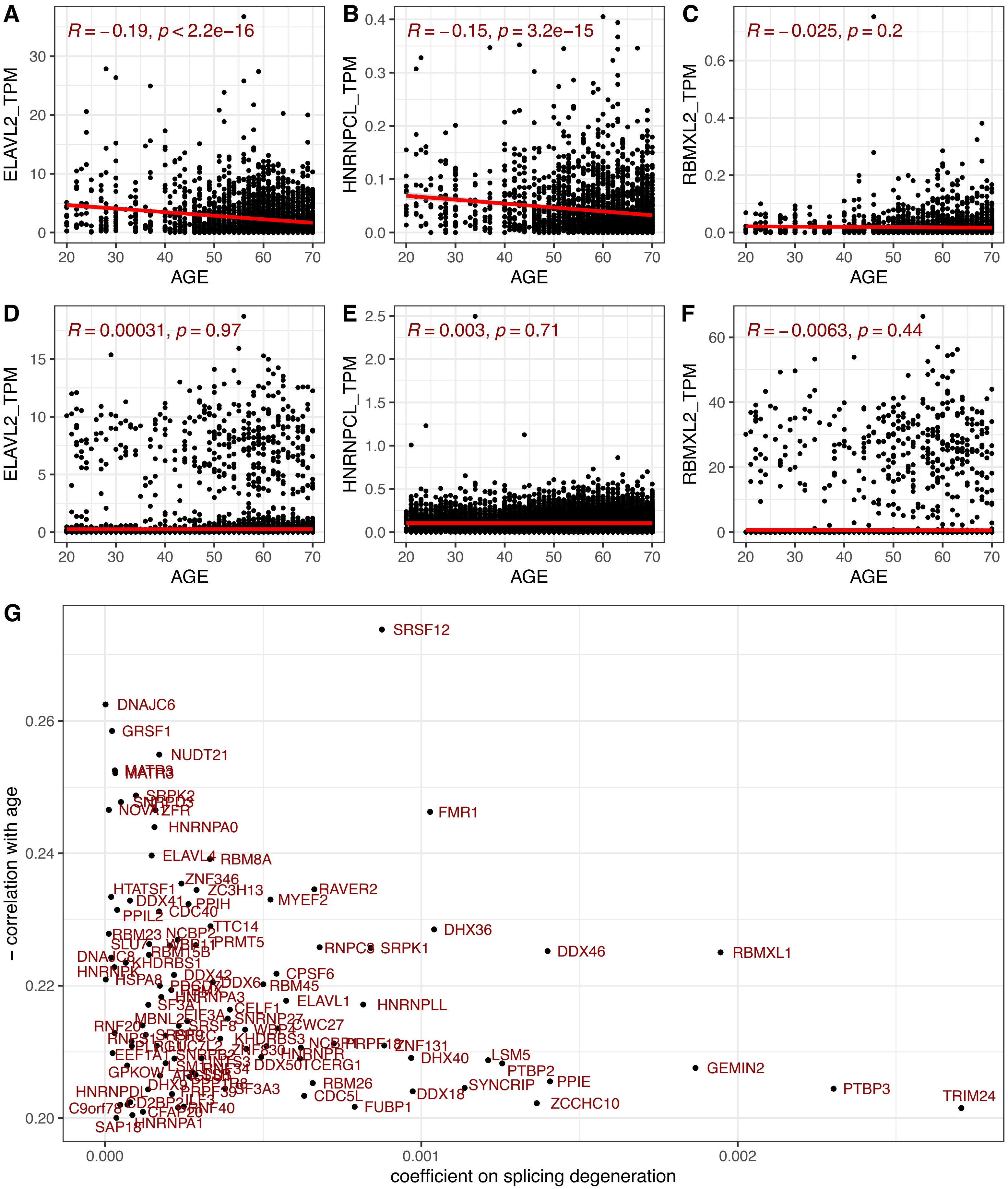
A
R = −0.19, p < 2.2e−16
ELAVL2_TPM
AGE
B
R = −0.15, p = 3.2e−15
HNRNPCL_TPM
AGE
C
R = −0.025, p = 0.2
RBMXL2_TPM
AGE
D
R = 0.00031, p = 0.97
ELAVL2_TPM
AGE
E
R = 0.003, p = 0.71
HNRNPCL_TPM
AGE
F
R = −0.0063, p = 0.44
RBMXL2_TPM
AGE
G
SRSF12
DNAJC6
GRSF1
NUDT21
FMR1
HNRNPA0
ELAVL4
RBM8A
ZNF346
HTATSF1
ZC3H13
RAVER2
MYEF2
DHX36
RNPC3
SRPK1
DDX46
RBMXL1
CPSF6
RBM45
ELAVL1
HNRNPLL
CWC27
ZNF131
DHX40
LSM5
PTBP2
GEMIN2
PPIE
SYNCRIP
RBM26
CDC5L
DDX18
FUBP1
ZCCHC10
PTBP3
TRIM24
SAP18
HNRNPA1
− correlation with age
coefficient on splicing degeneration

Figure S13

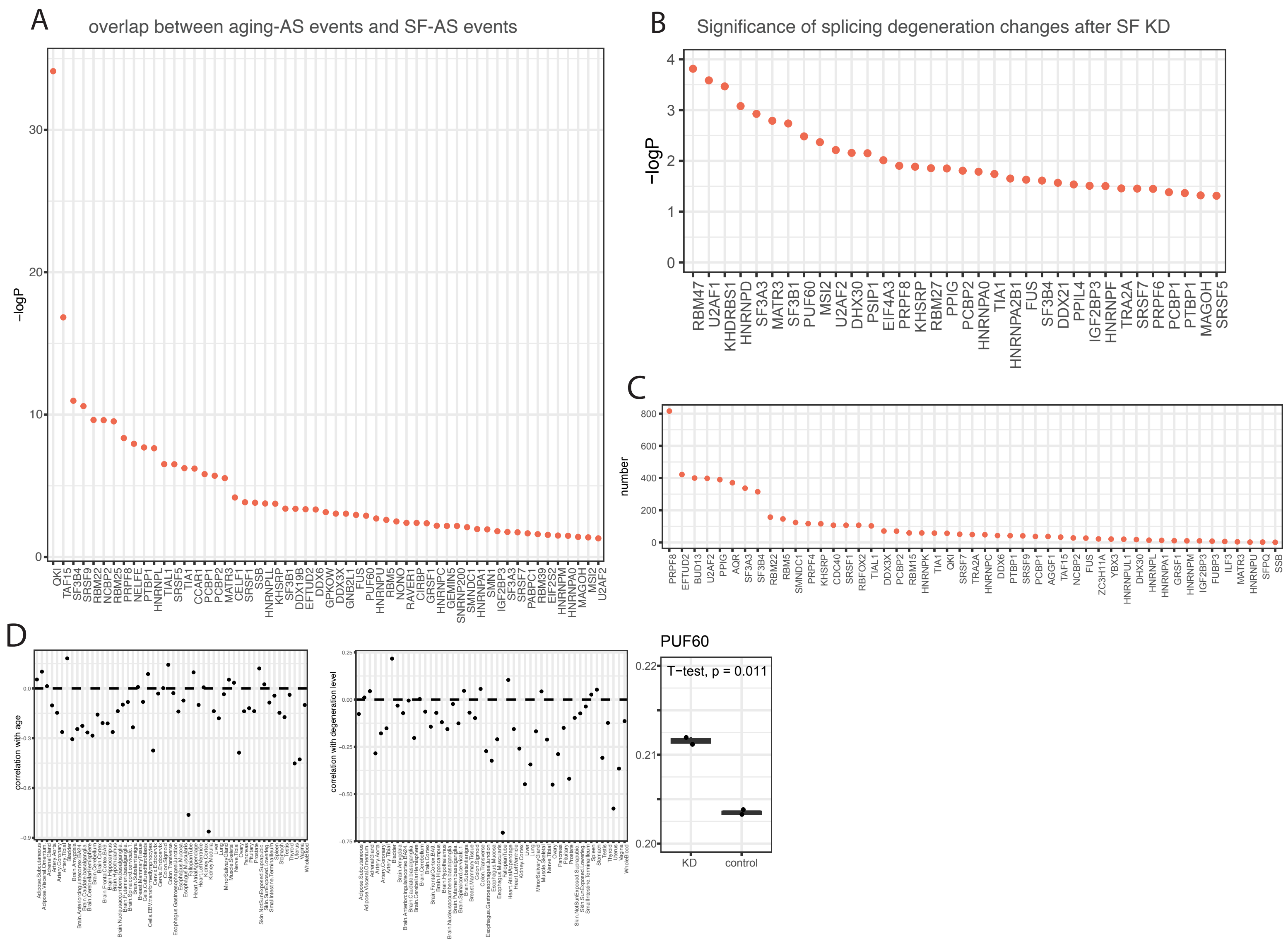
A
overlap between aging-AS events and SF-AS events
−logP
B
Significance of splicing degeneration changes after SF KD
−logP
C
number
D
correlation with age
correlation with degeneration level
PUF60
T−test, p = 0.011
KD
control

### Figure S14

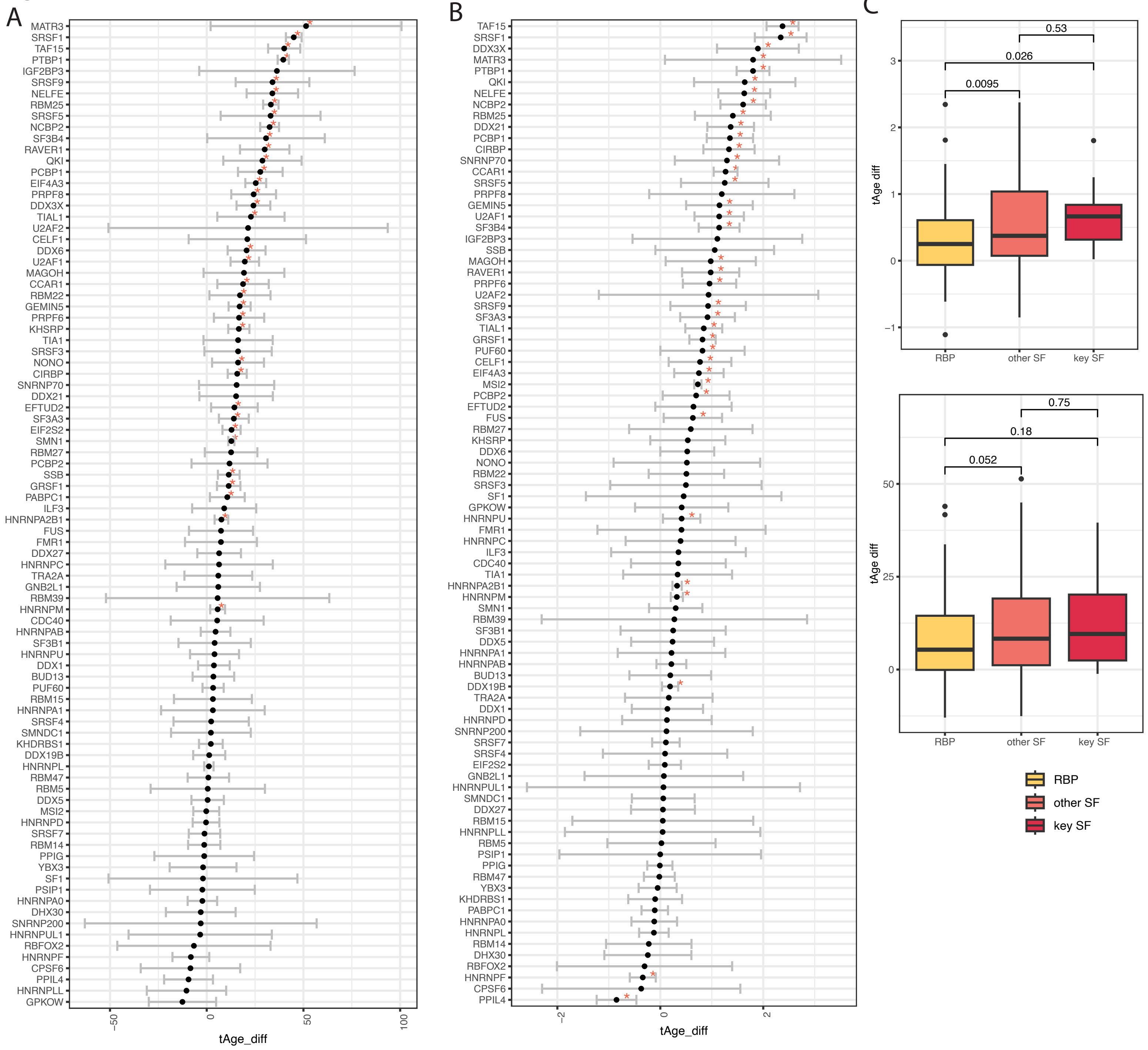

A
B
C
tAge_diff
tAge diff
RBP
other SF
key SF
0.53
0.026
0.0095
0.75
0.18
0.052

Figure S15

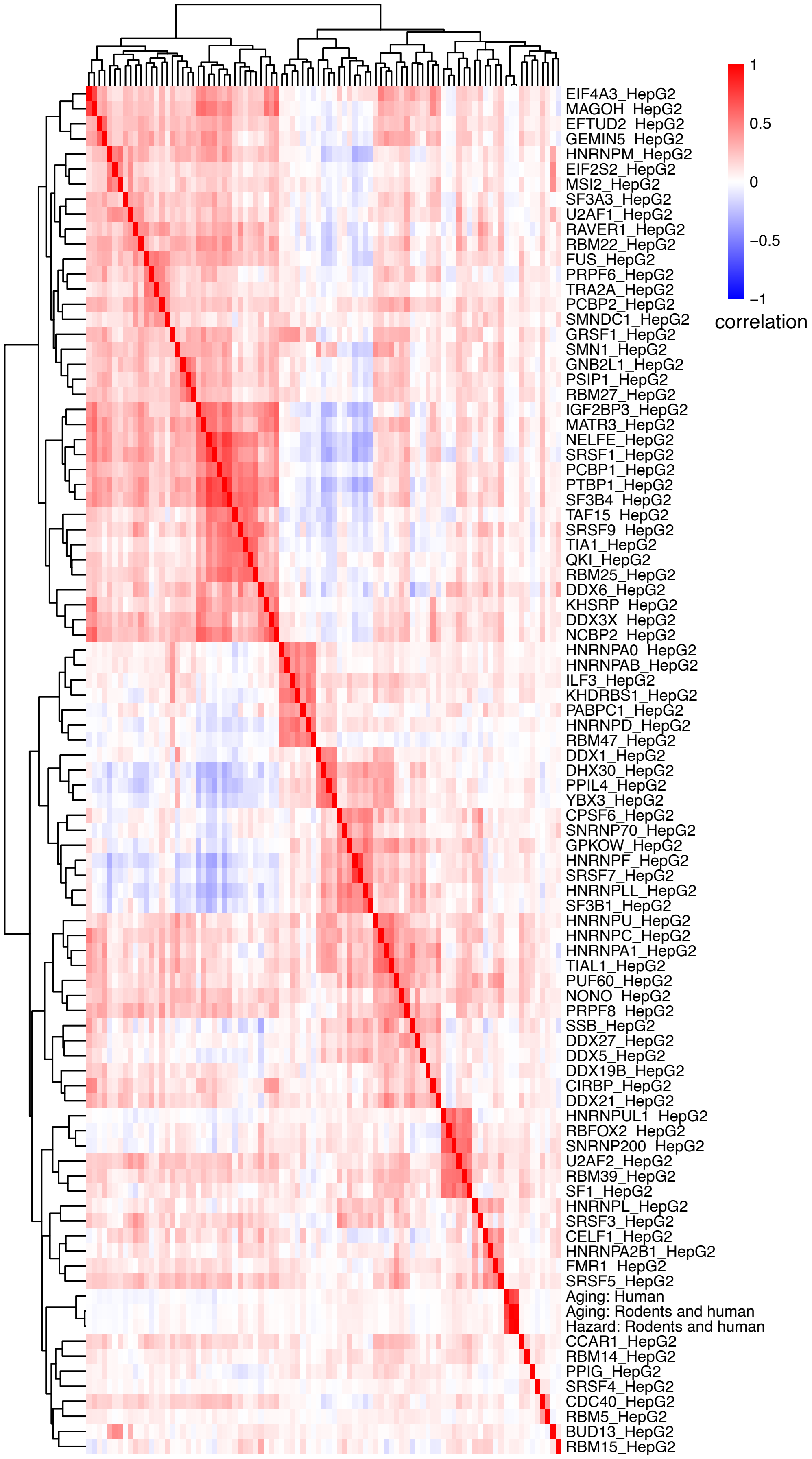
EIF4A3_HepG2
MAGOH_HepG2
EFTUD2_HepG2
GEMIN5_HepG2
HNRNPM_HepG2
EIF2S2_HepG2
MSI2_HepG2
SF3A3_HepG2
U2AF1_HepG2
RAVER1_HepG2
RBM22_HepG2
FUS_HepG2
PRPF6_HepG2
TRA2A_HepG2
PCBP2_HepG2
SMNDC1_HepG2
GRSF1_HepG2
SMN1_HepG2
GNB2L1_HepG2
PSIP1_HepG2
RBM27_HepG2
IGF2BP3_HepG2
MATR3_HepG2
NELFE_HepG2
SRSF1_HepG2
PCBP1_HepG2
PTBP1_HepG2
SF3B4_HepG2
TAF15_HepG2
SRSF9_HepG2
TIA1_HepG2
QKI_HepG2
RBM25_HepG2
DDX6_HepG2
KHSRP_HepG2
DDX3X_HepG2
NCBP2_HepG2
HNRNPA0_HepG2
HNRNPAB_HepG2
ILF3_HepG2
KHDRBS1_HepG2
PABPC1_HepG2
HNRNPD_HepG2
RBM47_HepG2
DDX1_HepG2
DHX30_HepG2
PPIL4_HepG2
YBX3_HepG2
CPSF6_HepG2
SNRNP70_HepG2
GPKOW_HepG2
HNRNPF_HepG2
SRSF7_HepG2
HNRNPLL_HepG2
SF3B1_HepG2
HNRNPU_HepG2
HNRNPC_HepG2
HNRNPA1_HepG2
TIAL1_HepG2
PUF60_HepG2
NONO_HepG2
PRPF8_HepG2
SSB_HepG2
DDX27_HepG2
DDX5_HepG2
DDX19B_HepG2
CIRBP_HepG2
DDX21_HepG2
HNRNPUL1_HepG2
RBFOX2_HepG2
SNRNP200_HepG2
U2AF2_HepG2
RBM39_HepG2
SF1_HepG2
HNRNPL_HepG2
SRSF3_HepG2
CELF1_HepG2
HNRNPA2B1_HepG2
FMR1_HepG2
SRSF5_HepG2
Aging: Human
Aging: Rodents and human
Hazard: Rodents and human
CCAR1_HepG2
RBM14_HepG2
PPIG_HepG2
SRSF4_HepG2
CDC40_HepG2
RBM5_HepG2
BUD13_HepG2
RBM15_HepG2
1
0.5
0
−0.5
−1
correlation
